# A multiple *Arc* (mArc) tagging system to uncover the organizational principles of multiple memories

**DOI:** 10.1101/2024.02.01.578410

**Authors:** Michelle Stackmann, Tushar D. Yelhekar, Meizhen Meng, Xiaochen Sun, Joseph Nthumba, Nicholas E. Bulthuis, Nick Vaughan, Elaine Zhu, Yingxi Lin, Christine A. Denny

## Abstract

Engrams or memory traces are the neuronal ensembles that collectively store individual experiences. Genetic strategies based on immediate early genes (IEGs), such as *Arc/Arg3.1*, allow us to tag the ensembles active during memory encoding and compare them to those active during retrieval. However, these strategies only allow for the tagging of one neural ensemble. Here, we developed a multiple *Arc* (mArc) system that allows for the tagging of two *Arc*^+^ ensembles. We validated this system by investigating how context, time, and valence influence neuronal ensemble reactivation in the dentate gyrus (DG). We show that similar contextual and valenced experiences are encoded in overlapping DG ensembles. We also find that ensembles are modulated by time, where experiences closer in time are encoded in more similar ensembles. These results highlight the dynamic nature of DG ensembles and show that the mArc system provides a powerful approach for investigating multiple memories in the brain.

**HIGHLIGHTS:**

- The mArc system allows for the tagging of two *Arc*^+^ ensembles in the same mouse
- DG ensembles labeled by the mArc system receive increased excitatory input
- Context, valence, and time influence DG ensemble reactivation
- DG neural ensembles are reactivated less with increasing time

## INTRODUCTION

The lasting physical and chemical changes in the brain due to learning constitute an engram or memory trace^1,2^. The cells that are active during learning, undergo plasticity changes due to the learning, and are necessary for memory retrieval are known as engram cells. These sparse, distributed set of cells are reactivated during memory retrieval^3,4^ and are necessary and sufficient for memory expression^4–9^. Although many studies have characterized the properties of single engrams in multiple conditions and various brain regions^8,10–12^, little is known about how multiple memories, and their respective ensembles, are co-stored in the brain.

Modern genetic tools have allowed for the identification and manipulation of the neural ensembles active during memory processes^3,4,6,13–16^. Many of these tools rely on immediate early genes (IEGs) such as *Arc*, *c-fos*, and *Npas4*, which are transiently and rapidly expressed in response to cellular stimuli^17^. In these systems, IEG promoters have been engineered to drive downstream expression of reporters (fluorescent, chemogenetic, or optogenetic) to label the cells active during particular memory processes. By tagging previously activated cells with a reporter, the cells active during memory encoding can be compared with those active during memory retrieval. A limitation of these strategies is that the labeling is confined to a single activated ensemble, thereby, prohibiting studies aimed at investigating multiple memories.

Although IEGs have been used interchangeably in engram studies, emerging evidence suggests that the ensembles defined by specific IEG expression can be functionally distinct components of memory. A recent study has shown that IEG-specific ensembles undergo different learning-induced synaptic changes and show opposing roles in memory-related behavior^18^, highlighting a critical need for investigating how IEG-specific ensembles participate in memory processes. The activity-regulated cytoskeleton-associated protein (*Arc*/*Arg3.1*) gene, unlike many IEGs that encode transcription factors, encodes an effector protein that directly regulates various forms of synaptic plasticity^19–24^. *Arc* is necessary for long-term memory consolidation^21,25–27^ and recent studies show that Arc proteins can form virus-like capsids that encapsulate and transfer mRNA from cell to cell^28,29^, giving *Arc* a potential role in intercellular communication. How *Arc* ensembles are activated during distinct memories is not known.

Here, we developed and characterized a novel multiple *Arc* (mArc) strategy to label two activated *Arc* ensembles in a single mouse. We show that the mArc strategy labels *Arc* ensembles that exhibit similar synaptic properties after a learning experience. We then used the mArc strategy to investigate how multiple contextual experiences are co-stored in the dentate gyrus (DG). We show that DG ensembles are reactivated in a context– and valence-specific manner, and that experiences closer in time are encoded in more overlapping DG ensembles. Together, these results show that the mArc strategy can be successfully used to uncover the organizational principles of multiple memories.

## RESULTS

### Low overlap between simultaneously tagged *Arc* and *c-fos* ensembles in the DG

To generate a system that labels multiple active ensembles in a single mouse, we first combined two activity-dependent labeling strategies **(Figure 1A)**. We used the ArcCreER^T2^ x enhanced yellow fluorescent protein (EYFP) transgenic line, in which an *Arc*^+^ ensemble can be permanently labeled with EYFP in a tamoxifen-dependent manner^4^. The other strategy we used is the well-characterized c-fos-tTA transgenic line, which allows for the long-lasting labeling of a *c-fos*^+^ ensemble under the control of doxycycline (DOX)^3^. In this line, the *c-fos* promoter drives expression of the tetracycline transactivator (tTA) that binds to the tetracycline-responsive element (TRE) to induce reporter expression. In the presence of DOX, tTA is prevented from binding to TRE, which then prevents reporter expression.

**Figure 1.**
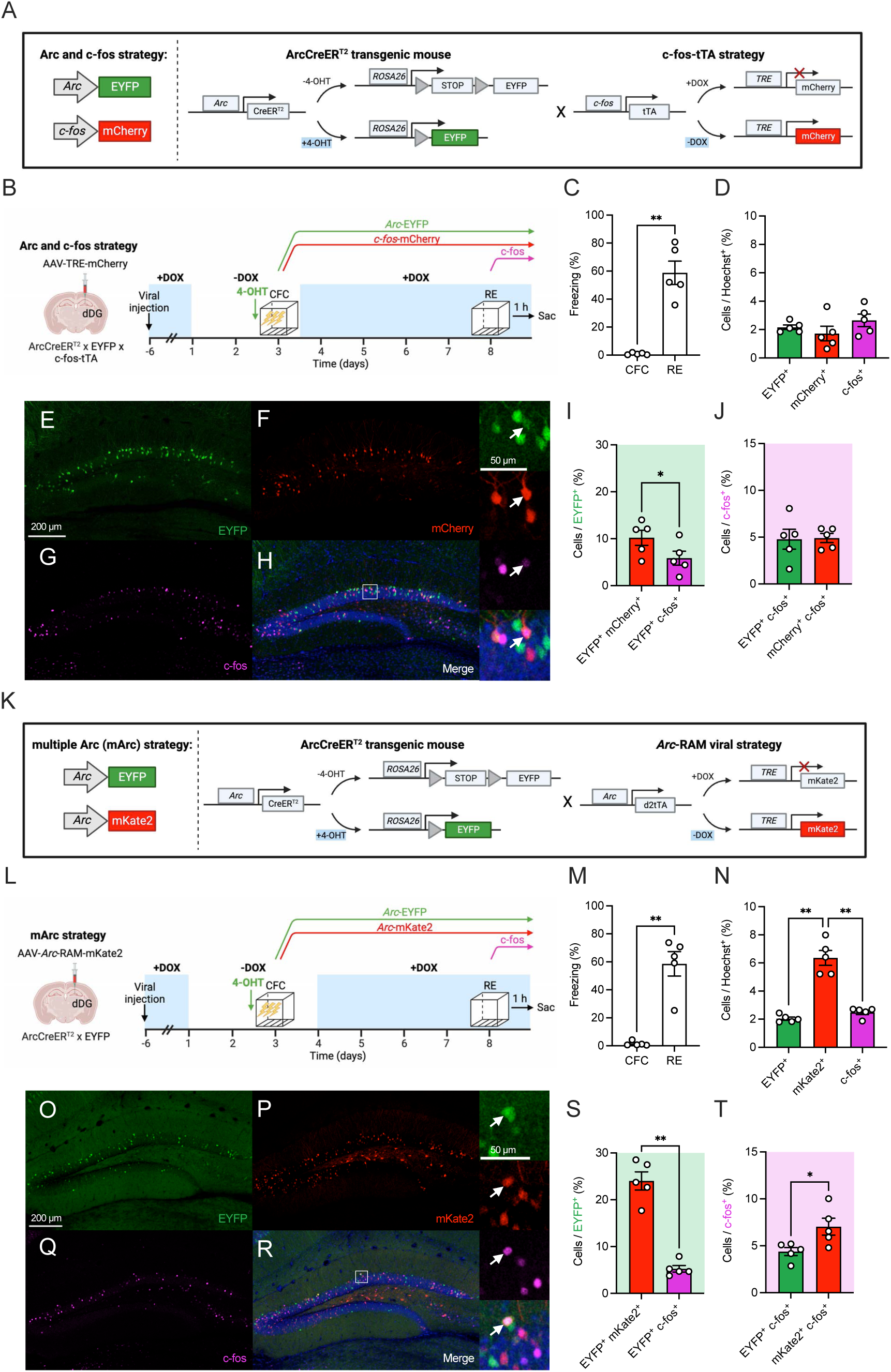
The mArc system labels overlapping *Arc* ensembles in the DG. (**A**) Genetic strategy to label an *Arc* and a *c-fos* ensemble in the same mouse using the ArcCreER^T^^2^ transgenic mouse line in combination with a c-fos-tTA strategy. (**B**) Experimental design to label an *Arc* and a *c-fos* fear ensemble at the same time in the DG. (**C**) Mice administered a 3-shock CFC paradigm freeze significantly more during re-exposure than during training. (**D**) The percentage of EYFP^+^, mCherry^+^, and c-fos^+^ cells is comparable. Representative images showing labeling of: (**E**) EYFP^+^ cells, (**F**) mCherry^+^ cells, (**G**) c-fos^+^ cells, and (**H**) merged DG section. The white box delineates the area magnified in the right panels. The arrows point to a triple-labeled cell. (**I**) The percentage of EYFP^+^ cells that also express mCherry is significantly greater than the percentage of EYFP^+^ cells that also express c-fos. (**J**) There is no difference in the percentage of c-fos^+^ cells that also express EYFP or mCherry. (**K**) Genetic strategy to label multiple *Arc* ensembles in the same mouse using the mArc system. (**L**) Experimental design to label a fear ensemble with the mArc strategy in the DG. (**M**) Mice administered a 3-shock CFC paradigm freeze significantly more during re-exposure than during training. (**N**) The percentage of mKate2^+^ cells is significantly higher than the percentage of EYFP^+^ and c– fos^+^ cells. Representative images showing labeling of: (**O**) EYFP^+^ cells, (**P**) mCherry^+^ cells, (**Q**) c-fos^+^ cells, and (**R**) Merged DG section. The white box delineates the area magnified in the right panels. The arrows point to a triple-labeled cell. (**S**) The percentage of EYFP^+^ cells that also express mKate2 is significantly greater than the percentage of EYFP^+^ cells that also express c-fos. **(T)** There is a higher percentage of c-fos^+^ cells that express mKate2 when compared to EYFP. n = 5 mice per group. Error bars show mean ± SEM. *p < 0.05, **p < 0.01. EYFP, enhanced yellow fluorescent protein; 4-OHT, 4-hydroxytamoxifen; tTA, tetracycline-controlled transactivator; DOX, doxycycline; TRE, tetracycline response element; dDG, dorsal dentate gyrus; CFC, contextual fear conditioning; RE, re-exposure; Sac, sacrifice; d2tTA, destabilized tetracycline transactivator; RAM, Robust Activity Marking.

To determine if the tagging strategies label comparable ensembles, we labeled the ensembles active during a single contextual fear conditioning (CFC) session with both strategies simultaneously **(Figure 1B)**. ArcCreER^T2^ x EYFP x c-fos-tTA mice were injected with an adeno-associated virus (AAV) encoding TRE-mCherry in the DG. Mice were taken off DOX and 2 days later injected with 4-hydroxytamoxifen (4-OHT) and subsequently subjected to CFC to label an *Arc* ensemble with EYFP and a *c-fos* ensemble with mCherry. After CFC, mice were placed back on DOX and 5 days later underwent re-exposure (RE) to the conditioned context and sacrificed 1 hour after. Mice froze upon RE to the conditioned context **(Figure 1C)**. The ensembles active during CFC were labeled with EYFP and mCherry, while the ensemble active during RE was visualized by staining for endogenous c-fos **(Figure 1E-1H)**. There was a similar percentage (∼2%) of labeled EYFP^+^, mCherry^+^, and c-fos^+^ cells in the granule cell layer of the DG **(Figure 1D)**. The percentage of cells labeled by both tagging strategies was relatively low, with only ∼10% of EYFP^+^ cells also being labeled by mCherry **(Figure 1I)**. However, this percentage was higher than that of the EYFP^+^ cells reactivated during memory retrieval (EYFP^+^ c-fos^+^ / EYFP^+^), showing that the two labeling strategies used simultaneously label more similar ensembles compared to the ensembles active during two fear experiences. Fear-encoding EYFP^+^ and mCherry^+^ ensembles were reactivated at a similar level during fear retrieval **(Figure 1J)**. Overall, the *Arc* and *c-fos* strategies labeled low overlapping ensembles, despite them labeling the same CFC experience.

To investigate if the low overlap between the ensembles tagged by the *Arc* and *c-fos* strategies was due to the combination of a transgenic and viral reporter, we developed fully transgenic strategy to label an *Arc* and a *c-fos* ensemble **(Figure S1A)**. We repeated the experiment as described in **Figure 1** using the quadruple transgenic ArcCreER^T2^ x EYFP x c-fos-tTA x tdTomato mice, with the exception that mice were taken off DOX for 2 or 3 days before CFC **(Figure S1B)**. Mice froze significantly during RE **(Figure S1C)**. There was reduced expression of tdTomato in the DG compared to EYFP and c-fos expression **(Figure S1D-S1O)**. The time off DOX did not affect tdTomato labeling; mice taken off DOX for 2 or 3 days showed similar tdTomato expression **(Figure S1Q)**. The number of EYFP^+^ cells remained consistent regardless of time off DOX, as expected **(Figure S1P)**. The overlap between the EYFP^+^ and tdTomato^+^ ensembles remained low for the quadruple transgenic strategy, with only ∼4% of EYFP^+^ cells expressing tdTomato and ∼13% of tdTomato^+^ cells expressing EYFP **(Figure S1R-S1S)**. We labeled the ensembles active during kainic acid administration, which globally increases hippocampal activity, to determine if both transgenic strategies were capable of labeling all DG cells **(Figure S1T)**. While there was robust labeling throughout the DG with the ArcCreER^T2^ x EYFP strategy, the c-fos x tdTomato strategy incompletely labeled DG cells **(Figure S1U-S1X)**. Overall, these data show that the quadruple strategy incompletely labels an active *c-fos*^+^ ensemble.

### Development of a novel *Arc*-RAM viral strategy

We reasoned that the low overlap between the labeled *Arc* and *c-fos* labeling strategies could also be due to the different IEG promoters, and that using labeling systems with the same IEG promoter would lead to a higher overlap between labeled ensembles. Therefore, we developed a viral strategy based on the Robust Activity Marking (RAM) system^14,18^ that allows for the long-lasting tagging of *Arc^+^* ensembles **(Figure S2A)**. This strategy, which we called *Arc*-RAM, labels ensembles in a DOX-dependent manner. The promoter used in *Arc*-RAM is a short, synthetic promoter based on elements of the *Arc* enhancer/promoter region that regulate *Arc* induction, called the enhanced synaptic activity-responsive element (E-SARE)^30^.

To characterize *Arc*-RAM *in vivo*, 129S6/SvEv mice were injected with *Arc*-RAM into the DG and the ensembles active in the home cage (HC, OFF DOX), during CFC training (CFC, OFF DOX), or on DOX (CFC, ON DOX) were labeled **(Figure S2B)**. For the OFF DOX groups, mice were kept on DOX until 48 h before behavior and for the next 24 h, as previously described^14^. The percentage of labeled mKate2^+^ cells in the home cage was lower than the percentage labeled during CFC (∼2% vs. 4%, respectively) **(Figures S2C-S2D, S2F)**. The CFC, ON DOX mice showed minimal mKate2 labeling compared to CFC, OFF DOX mice **(Figure S2E-S2F)**. These data indicate that *Arc*-RAM effectively labels active ensembles and is tightly regulated by DOX.

### The mArc system labels highly overlapping ensembles in the DG

To test if labeling strategies driven by the same IEG promoter would lead to more similarly labeled ensembles, we created a multiple *Arc* (mArc) system by combining the ArcCreER^T2^ and *Arc*-RAM strategies **(Figure 1K).** Tagging of *Arc* ensembles simultaneously occurred as described in **Figure 1B (Figure 1L).** Mice froze during significantly during RE **(Figure 1M)**. The ensembles active during CFC were labeled with EYFP and mKate2, and the ensemble active during RE was visualized by staining for endogenous c-fos **(Figure 1O-1R)**. There was a higher percentage of labeled mKate2^+^ cells than EYFP^+^ or c-fos^+^ cells (∼6% vs 2%) **(Figure 1N)**. The percentage of cells labeled by two *Arc* tagging strategies was higher than that of an *Arc* and a *c-fos* tagging strategy, with ∼25% of EYFP^+^ cells also expressing mKate2 **(Figure 1S)**. This percentage was higher than the percentage of EYFP^+^ cells reactivated during memory retrieval (EYFP^+^ c-fos^+^ / EYFP^+^). A higher percentage of fear-encoding mKate2^+^ cells was reactivated during retrieval compared to EYFP^+^ cells (∼4% vs. 7%) **(Figure 1T)**, which is likely due to a greater number of cells being labeled with mKate2 compared to EYFP. Overall, the mArc system labels more overlapping ensembles than an *Arc* and *c-fos* labeling strategy.

### Maximum overlap between tagging strategies for the mArc system in the DG

To investigate the maximum overlap between the ensembles tagged by the mArc system, we labeled the cells active during a kainic acid-induced seizure with the two tagging strategies simultaneously **(Figure 2A)**. A kainic acid-induced seizure resulted in EYFP and mKate2 labeling throughout the DG, showing robust activity-dependent labeling for both *Arc* strategies **(Figure 2D-2L)**. The percentage of EYFP^+^ cells that also expressed mKate2 was ∼45%, showing a substantial co-localization between the tagging strategies **(Figure 2B)**. A lower percentage of mKate2^+^ cells also expressed EYFP (i.e., 23%) **(Figure 2C)**. The incomplete overlap between strategies could be due to promoter competition, in which one strategy/reporter is selectively expressed within a cell due to the preferential binding to that particular strategy’s promoter. We investigated if a reduced viral load would lead to a higher overlap between the labeled ensembles but observed that reducing the *Arc*-RAM viral load led to incomplete viral expression in the DG **(Figure S3)**. These results establish a value of maximum overlap between the tagging strategies (∼45%) and show that, despite incomplete co-expression, both strategies are robustly induced by activity in the DG.

**Figure 2.**
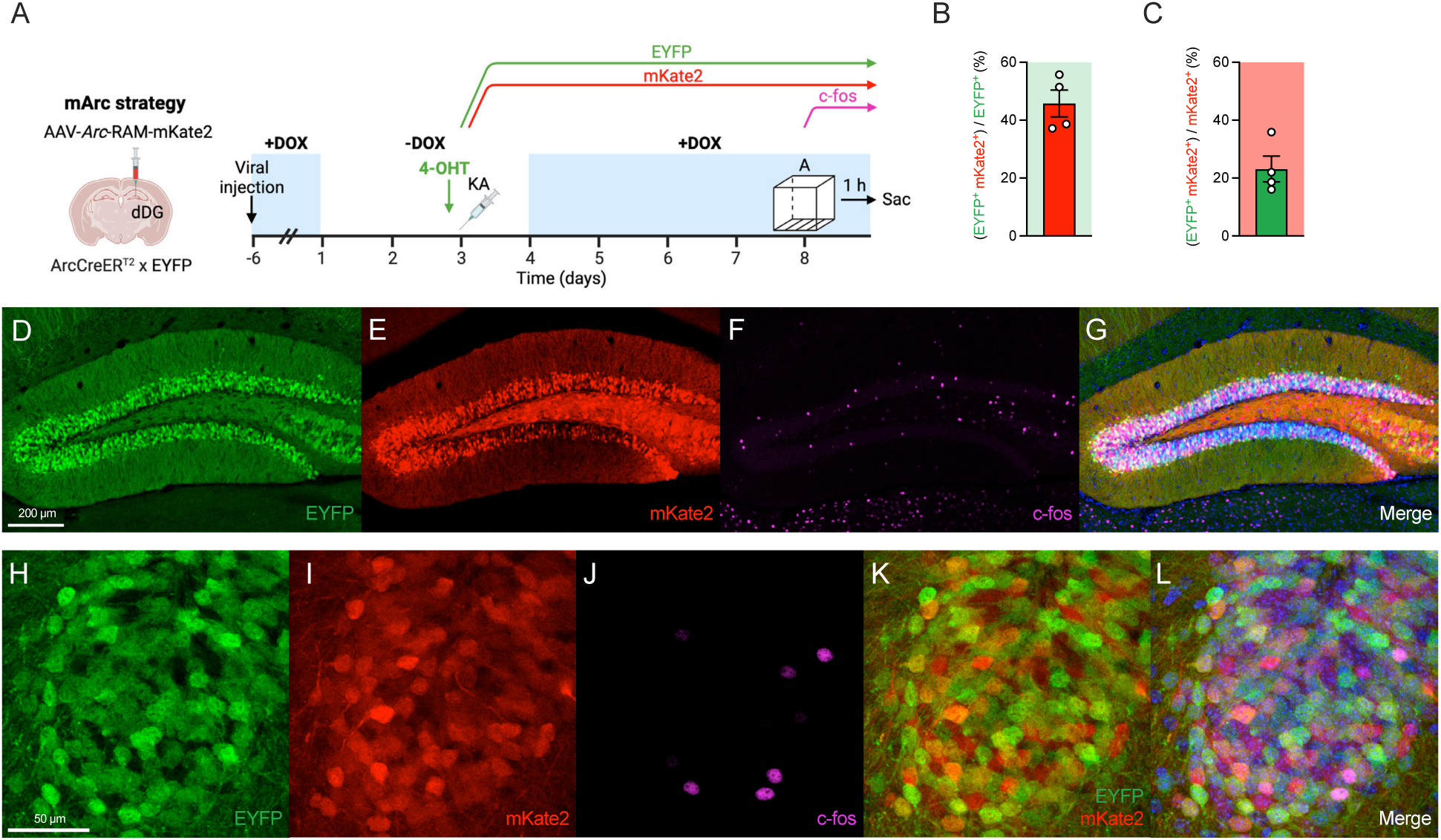
mArc system labeling in the DG is robustly induced by kainic acid-induced seizure. (**A**) Experimental design to label active ensembles during seizure with the mArc system. (**B**) Approximately 45% of EYFP^+^ cells are co-labeled with mKate2. (**C**) Approximately 23% of mKate2^+^ cells are co-labeled with EYFP. Representative images of: (**D**) EYFP^+^ cells, (**E**) mKate2^+^ cells, (**F**) c-fos^+^ cells, and (**G**) merged DG section. Representative magnified images of: (**H**) EYFP^+^ cells, (**I**) mKate2^+^ cells, (**J**) c-fos^+^ cells, (**K**) EYFP^+^ and mKate2^+^ cells, and (**L**) merged DG section. n = 4 mice. Error bars show mean ± SEM. EYFP, enhanced yellow fluorescent protein; RAM, Robust Activity Marking; dDG, dorsal dentate gyrus; DOX, doxycycline; 4-OHT, 4-hydroxytamoxifen; KA, kainic acid; Sac, sacrifice.

### The mArc system labels ensembles with similar synaptic properties in the DG

To further validate that the mArc system labels similar ensembles, we investigated the electrophysiological properties of tagged *Arc^+^*ensembles. ArcCreER^T2^ mice were injected with *Arc*-RAM and AAV5-DIO-EYFP into the DG **(Figure 3A)**. Mice were taken off DOX and 2 days later injected with 4-OHT and subsequently subjected to CFC to label fear-encoding *Arc* ensembles with EYFP and mKate2. Five to seven days later, whole-cell patch clamp recordings of EYFP^+^, mKate2^+^, double-labeled EYFP^+^ mKate2^+^, and unlabeled neighboring granule cells were performed. Labeled EYFP^+^, mKate2^+^, and EYFP^+^ mKate2^+^ cells showed an increase in miniature excitatory postsynaptic current (mEPSC) frequency compared to unlabeled cells, indicating that labeled cells receive an increased number of excitatory inputs **(Figure 3B-3C)**. There was no difference in the mEPSC amplitude between labeled and unlabeled cells **(Figure 3D)**. There was no difference in passive membrane properties between EYFP^+^, mKate2^+^, double-labeled EYFP^+^ mKate2^+^, and unlabeled cells **(Figure 3E-3M)**. These data show that the ensembles labeled by both *Arc* labeling strategies are similar in terms of their synaptic properties and receive an increased number of excitatory inputs compared to unlabeled cells.

**Figure 3.**
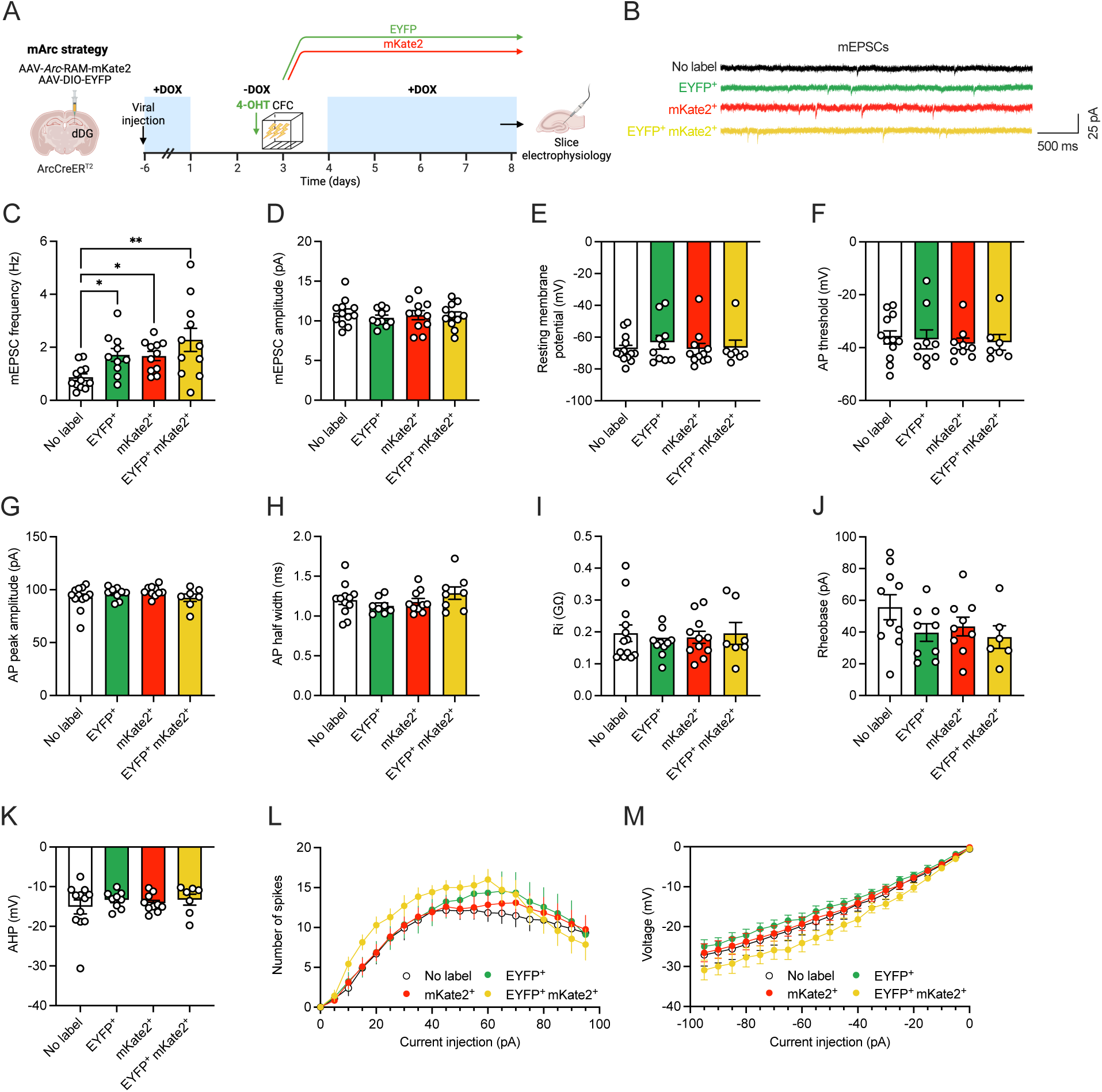
DG ensembles labeled by the mArc system receive increased excitatory input. (**A**) Experimental design to label a fear ensemble in the DG with the mArc system and measure the synaptic properties of EYFP^+^, mKate2^+^, EYFP^+^ mKate2^+^, and unlabeled neurons. (**B**) Representative traces showing recorded mEPSCs from EYFP^+^, mKate2^+^, EYFP^+^ mKate2^+^, and unlabeled neurons. (**C**) EYFP^+^, mKate2^+^, and EYFP^+^ mKate2^+^ neurons show increased mEPSC frequency when compared to unlabeled neurons. (**D**) There is no difference in the mEPSC amplitude of EYFP^+^, mKate2^+^, EYFP^+^ mKate2^+^ and unlabeled neurons. There is no difference in the: (**E**) resting membrane potential, (**F**) AP threshold, (**G**) AP peak amplitude, (**H**) AP half width, (**I**) input resistance (Ri), (**J**) rheobase, and (**K**) afterhyperpolarization (AHP) of EYFP^+^, mKate2^+^, EYFP^+^ mKate2^+^ and unlabeled neurons. (**L**) Current injection vs. number of spikes recorded from EYFP^+^, mKate2^+^, EYFP^+^ mKate2^+^, and unlabeled neurons. There is no difference in number of spikes between groups. (**M**) Current injection vs. voltage response recorded form EYFP^+^, mKate2^+^, EYFP^+^ mKate2^+^, and unlabeled neurons. There is no difference in the voltage responses between groups. n = 6-13 neurons per group. Error bars show mean ± SEM. *p < 0.05, **p < 0.01. EYFP, enhanced yellow fluorescent protein; dDG, dorsal dentate gyrus; DOX, doxycycline; 4-OHT, 4-hydroxytamoxifen; CFC, contextual fear conditioning.

### The mArc system labels ensembles consistent with the cell-type specificity of *Arc* expression

*Arc* is exclusively expressed in neurons and co-localizes with excitatory neurons in the hippocampus^31^. To confirm the cell-type specificity of the mArc system labeling in the DG, we analyzed the co-localization of mArc labeled cells with parvalbumin (PV), a marker of fast-spiking interneurons^32^ **(Figure S4A-S4G)**, and with neuronal nuclear protein (NeuN), a neuronal marker^33^ **(Figure S4H-S4N)**. We observed that EYFP^+^ and mKate2^+^ ensembles did not co-localize with PV^+^ cells and always co-localize with NeuN. Our results show the mArc system labels ensembles consistent with the cell-type specificity of *Arc* expression in the hippocampus.

### Context and time modulate ensemble reactivation in the DG

Following validation of the mArc system, we first sought to determine how multiple contextual experiences are represented at the ensemble level in the DG. Here, we used the mArc strategy to label the ensembles active during multiple exposures to the same or different contexts **(Figure 4A)**. ArcCreER^T2^ x EYFP mice were injected with *Arc*-RAM. Mice were injected with 4-OHT and subsequently exposed to context A to label active neurons with EYFP. Five days later, after mice had been taken off DOX for 2 days, one group of mice was exposed to context A (Same Context group) and another group was exposed to a slightly different context B (Different Context group) to label active neurons with mKate2. One day later, the Same Context group was exposed to context A, and the Different Context group was exposed to a distinct context C and sacrificed 1 hour later **(Figure 4C)**. Mice in the Same Context group showed increased immobility during the second and third exposures to context A compared to the first exposure, indicating habituation to the familiar context **(Figure 4B)**. Mice in the Different Context group showed no difference in immobility levels during all context exposures. There was a similar percentage of EYFP^+^ and mKate2^+^ cells labeled between the two groups **(Figure 4D-4E)**. A greater percentage of c-fos^+^ cells was labeled in the Same Context group when compared to the Different Context group **(Figure 4F)**.

**Figure 4.**
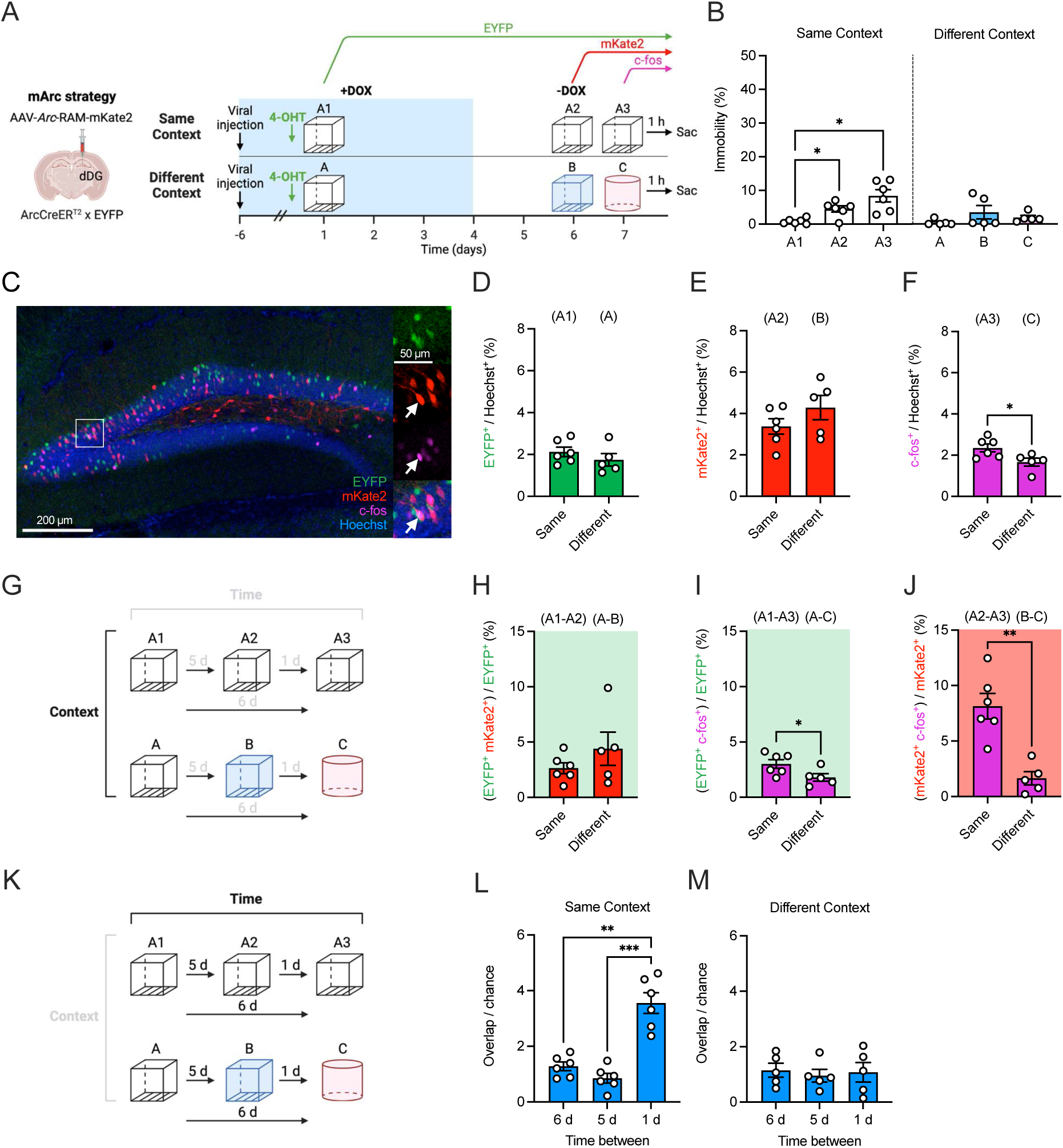
Context and time modulate ensemble reactivation in the DG. (**A**) Experimental design to label the ensembles active during three exposures to the same context or three exposures to different contexts. (**B**) Mice in the Same Context group show increased immobility in the second and third exposures to context A compared to the first exposure. Mice in the Different Context group show no difference in immobility during exposures to the different contexts. (**C**) Representative image of EYFP^+^, mKate2^+^ and c-fos^+^ ensembles in the dDG. The white box delineates the area magnified in the right panels. The white arrows point to a mKate2^+^ c-fos^+^ cell. (**D**) The percentage of EYFP^+^ cells does not differ between the Same Context and Different Context groups. (**E**) The percentage of mKate2^+^ cells does not differ between the Same Context and Different Context groups. (**F**) The Same Context group shows a greater percentage of c-fos^+^ cells compared to the Different Context group. (**G**) Diagram showing between-group comparisons. (**H**) The percentage of EYFP^+^ cells that also express mKate2 does not differ between the Same Context and Different Context groups. (**I**) The Same Context group has a greater percentage of EYFP^+^ cells that also express c-fos when compared to the Different Context group. (**J**) The Same Context group has a greater percentage of mKate2^+^ cells that also express c-fos compared to the Different Context group. (**K**) Diagram showing within-group comparisons. (**L**) The overlap between ensembles active 1 d apart is greater than that of ensembles active 6 d or 5 d apart for the Same Context group. (**M**) There is no difference in the overlap between ensembles active 6, 5, or 1 d apart for the Different Context group. n = 5-6 mice per group. Error bars show mean ± SEM. *p < 0.05, **p < 0.01, ***p < 0.001. dDG, dorsal dentate gyrus; EYFP, enhanced yellow fluorescent protein; 4-OHT, 4-hydroxytamoxifen; DOX, doxycycline; Sac, sacrifice.

To investigate the effect of context on the reactivation of ensembles, we first compared the reactivation of ensembles in a between-group manner **(Figure 4G)**. We compared the percentage of reactivated cells between the first and second context exposures. There was no difference in the percentage of (EYFP^+^ mKate2^+^) / EYFP^+^ cells between the Same Context group and Different Context group **(Figure 4H)**. We then compared the percentage of reactivated cells between the first and third context exposures. A greater percentage of (EYFP^+^ c-fos^+^) / EYFP^+^ cells was observed in the Same Context group when compared to the Different Context group (3% vs. 1.8%) **(Figure 4I)**. We then compared the percentage of reactivated cells between the second and third context exposures. A greater percentage of (mKate2^+^ c-fos^+^) / mKate2^+^ cells was observed in the Same Context group when compared to the Different Context group (8.1% vs. 1.6%) **(Figure 4J)**. These results indicate that DG ensembles are reactivated in a context-specific manner across multiple contextual exposures.

Apart from allowing us to investigate how multiple ensembles are reactivated between groups, the mArc system allows us to assess the effects of multiple time intervals on ensemble reactivation by comparing how ensembles are reactivated in a within-group manner **(Figure 4K)**. Due to differences in the percentage of cells labeled by each tagging system, the reactivation rates between ensembles were normalized. For each pair of labeled ensembles, we divided the percentage of observed co-labeled cells by the percentage of cells we would expect to see co-labeled due to chance. For the Same Context group, the overlap between the ensembles active 1 d apart was greater than that of the ensembles active 5 or 6 days apart **(Figure 4L)**. In contrast, there was no difference in the overlap of the ensembles active 6, 5, or 1 day(s) apart for the Different Context group **(Figure 4M)**. Similar results were obtained when Arc was used to label the third active ensemble instead of c-fos **(Figure S5A-S5E)**. Together, these data show that DG ensembles are reactivated in a context-specific manner and that reactivation is modulated by the time between experiences.

To investigate if the number of experiences or the time between experiences is driving an increase in reactivation, we labeled the ensembles active during two exposures to the same context, with one day in between, and quantified the overlap between the ensembles **(Figure S5F-S5H)**. We also labeled the ensembles active during three exposures to the same context, with five days in between experiences, and quantified the overlap between ensembles **(Figure S5I-S5M)**. We observed that the increase in reactivation was due to the short time between experiences and not the number of experiences.

### Ensemble reactivation in CA3 is not modulated by time

To test the versatility of the mArc system in additional brain regions, we characterized how multiple contextual experiences are encoded in hippocampal CA3. ArcCreER^T2^ x EYFP mice were injected with *Arc*-RAM into CA3 and three exposures to context A were tagged with EYFP, mKate2, and c-fos as shown in **Figure 4 (Figure S6A, S6C-S6F)**. Mice showed increased immobility during the second and third context exposures compared to the first exposure, indicating habituation **(Figure S6B)**. The percentage of labeled cells increased with the number of context exposures (1.4%, 6.3%, and 23% of cells) **(Figure S6G)**. Interestingly, when comparing the reactivation of ensembles between exposures, the reactivation between the first and third exposure **(Figure S6I)** and second and third exposure **(Figure S6J)** was greater in CA3 than in the DG **(Figure S6H-S6J)**, potentially due to the high percentage of CA3 cells active during the third exposure and the role of CA3 in pattern completion^34^. Unlike in the DG, there was no significant difference in the overlap of ensembles active 6, 5, or 1 day(s) apart in CA3 **(Figure S6K)**. These results show that the mArc strategy is successful in labeling ensembles other brain regions and that contextual ensembles are modulated differently within hippocampal regions.

### Fear ensemble reactivation is similar regardless of fear training strength

To determine how fear-encoding ensembles are reactivated during multiple fear experiences, with or without additional fear training, we used the mArc strategy to label the ensembles active during 1– or 2-trial CFC and subsequent re-exposures to the fear context **(Figure 5A)**. Mice underwent 3-shock CFC and active neurons were labeled with EYFP. Five days later, one group of mice was re-exposed to the conditioned context (1-Trial CFC group) and another group was administered 3-shock CFC again (2-Trial CFC group), and the active neurons were labeled with mKate2. One day later, mice from both groups were re-exposed to the conditioned context and sacrificed 1 hour later **(Figure 5C)**. In the 1-Trial CFC group, mice froze during the first and second re-exposure **(Figure 5B)**. In the 2-Trial CFC group, mice froze significantly during the second conditioning session and final re-exposure. Mice in the 2-Trial CFC group showed higher freezing levels in the last re-exposure when compared to the mice in the 1-Trial CFC group, showing a relative strengthening of the fear memory. There was a similar percentage of EYFP^+^, mKate2^+^, and c-fos^+^ cells labeled between groups, regardless of whether the conditioning response was strengthened or not **(Figure 5D-5F)**. To investigate the effect of memory strength on ensemble reactivation, we compared the reactivation of ensembles in a between-group manner **(Figure 5G)**. We first compared the percentage of reactivated cells between the first and second exposures, which were CFC and RE for the 1-Trial CFC group and two CFC experiences for the 2-Trial CFC group. There was no difference in the percentage of (EYFP^+^ mKate2^+^) / EYFP^+^ cells between the groups **(Figure 5H)**. We then compared the percentage of reactivated cells between the first and third exposures, which were CFC and a second RE for the 1-Trial CFC group and CFC and final RE for the 2-Trial CFC group. There was no difference in the percentage of (EYFP^+^ c-fos^+^) / EYFP^+^ cells between the groups **(Figure 5I)**. Finally, we compared the percentage of reactivated cells between the second and third exposures, which were two RE for the 1-Trial CFC group and a second CFC experience and RE for the 2-Trial CFC group. There was no difference in the percentage of (mKate2^+^ c-fos^+^) / mKate2^+^ cells between the groups **(Figure 5J)**. These results show that DG ensembles are reactivated similarly regardless of the strength of the memory.

**Figure 5.**
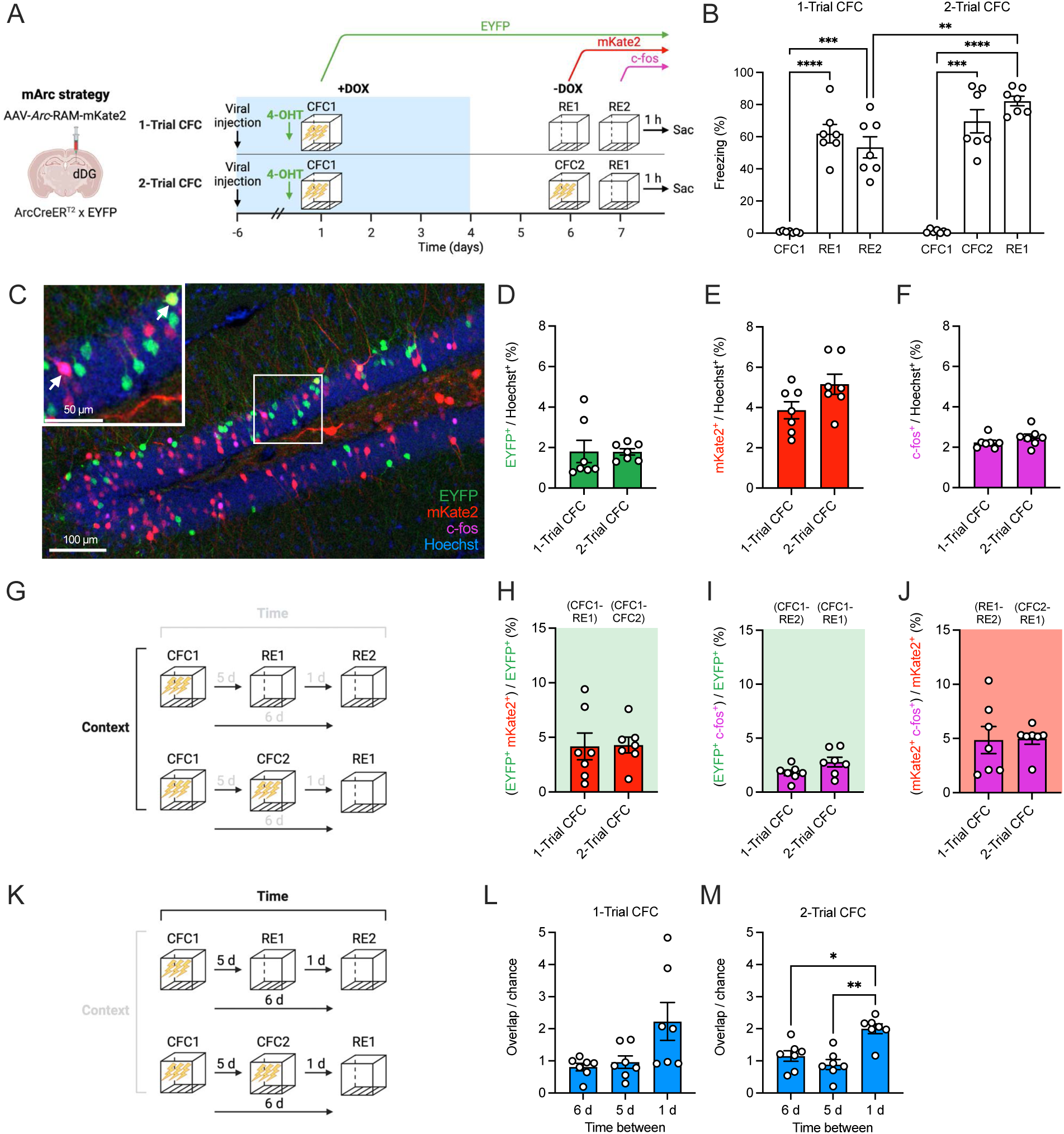
Fear ensemble reactivation is similar regardless of fear training strength. (**A**) Experimental design to label the ensembles active during 1– or 2-trial CFC and subsequent re-exposure to the fearful context. (**B**) Mice in the 1– and 2-Trial CFC groups freeze during re-exposures to the fearful context. In the last context re-exposure, mice in the 1-Trial CFC group show decreased freezing compared to the mice in the 2-Trial CFC group. (**C**) Representative image of EYFP^+^, mKate2^+^ and c-fos^+^ ensembles in the dDG. The white box delineates the area magnified in the upper left panel. The arrows point to double-labeled cells. The percentage of: (**D**) EYFP^+^ cells, (**E**) mKate2^+^ cells, and (**F**) c-fos^+^ cells do not differ between the 1-Trial CFC and 2-Trial CFC groups. (**G**) Diagram showing between-group comparisons. (**H**) The percentage of EYFP^+^ cells that also express mKate2 does not differ between groups. (**I**) The percentage of EYFP^+^ cells that also express c-fos does not differ between groups. (**J**) The percentage of mKate2^+^ cells that also express c-fos does not differ between groups. (**K**) Diagram showing within-group comparisons. (**L**) There is a trending but non-significant difference in the overlap between ensembles active 6, 5, or 1 d apart for the 1-Trial CFC group. (**M**) The overlap between ensembles active 1 d apart is greater than that of ensembles active 6 or 5 d apart in the 2-Trial CFC group. n = 7 mice per group. Error bars show mean ± SEM. *p < 0.05, **p < 0.01, ***p < 0.001, ****p < 0.0001. dDG, dorsal dentate gyrus; EYFP, enhanced yellow fluorescent protein; 4-OHT, 4-hydroxytamoxifen; DOX, doxycycline; CFC, contextual fear conditioning; RE, re-exposure; Sac, sacrifice.

To assess how fear-encoding ensembles are reactivated across time, we normalized the reactivation rates between pairs of experiences within groups **(Figure 5K)**. For the 1-Trial CFC group, there was a trending but non-significant difference between the reactivation rates of ensembles 6, 5, or 1 day(s) apart (repeated measures one-way ANOVA, p = 0.099) **(Figure 5L)**. For the 2-Trial CFC group, the reactivation of the ensembles 1 day apart was greater than that of the ensembles 6 or 5 days apart **(Figure 5M)**. These data show that DG fear ensemble reactivation is modulated by time, with experiences closer in time being encoded in more overlapping ensembles.

### Positive and negative ensembles are reactivated during re-exposure to a valenced experience

To determine how distinctly valenced experiences are represented at the ensemble level in the DG, we used the mArc system to label the ensembles active during a negative experience (i.e., CFC), a positive experience (i.e., social investigation), and re-exposure to either the negative or positive condition **(Figure 6A)**. Mice underwent 3-shock CFC and active neurons were labeled with EYFP. Five days later, mice were exposed to a female mouse in a distinct context and active neurons were labeled with mKate2. One day later, mice were again administered CFC (Fear retrieval group) or were re-exposed to the female (Social retrieval group) and sacrificed 1 hour later. Mice in the Fear retrieval group froze upon re-exposure to the CFC context **(Figure 6B-6C)**. To quantify memory of the female stimulus, mice were able to freely investigate the female under a cup or an empty cup. The time spent investigating the female mouse and the empty cup was quantified to assess memory. In the first exposure to females, mice spent more time investigating the female than the empty cup **(Figure 6D, 6F)**. In the second exposure to females, mice spent a similar amount of time investigating the female and the empty cup, showing recognition of the female mouse **(Figure 6E-6F)**. Mice also traveled less distance during the second female exposure than the first time, showing habituation to the context **(Figure 6G)**. The percentage of EYFP, mKate2, and c-fos cells was quantified **(Figure 6H)**. The percentage of mKate2^+^ cells was greater than EYFP^+^ cells **(Figure 6I)**. There was no difference in the number of c-fos^+^ cells active in either group **(Figure 6J)**. To investigate the effect of valenced experiences on ensemble reactivation, we compared ensemble reactivation between groups **(Figure 6K)**. We compared the percentage of reactivated cells between the first fear and social experiences. There was no difference in the percentage of (EYFP^+^ mKate2 ^+^) / EYFP^+^ cells between the groups **(Figure 6L)**. We then compared the percentage of reactivated cells between the first and third exposures; a greater percentage of (EYFP^+^ c-fos^+^) / EYFP^+^ cells was reactivated in the Fear retrieval group (which corresponded to two fear experiences) when compared with the Social retrieval group (which corresponded to a fear and a social experience) (2% vs. 1%) **(Figure 6M)**. We then compared the percentage of reactivated cells between the second and third exposures, which were a social experience and re-exposure to the fear context (Fear retrieval group) or social experience and re-exposure to the social context (Social retrieval group). A greater percentage of (mKate2^+^ c-fos^+^) / mKate2^+^ cells were reactivated in the Social retrieval group when compared with the Fear retrieval group (6% vs. 4%) **(Figure 6N)**. Overall, these results show that positive and negative ensembles are reactivated in a valence-specific manner.

**Figure 6.**
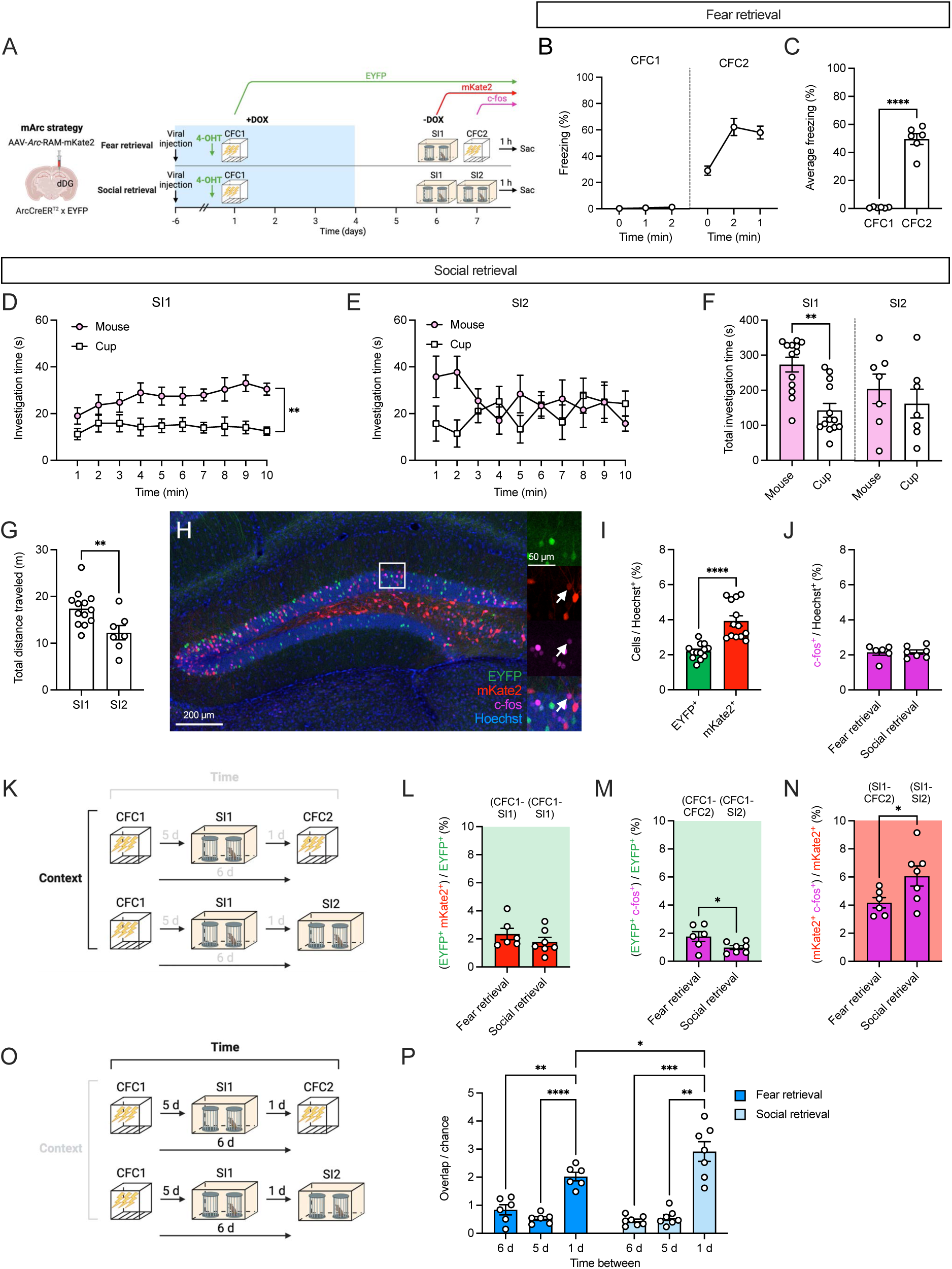
Positive and negative ensembles are reactivated during re-exposure to a valenced experience. (**A**) Experimental design to label the ensembles active during either positive (i.e., social) or negative (i.e., fear conditioning) experiences. Mice underwent CFC, social interaction, and then an additional CFC experience (Fear retrieval) or an additional social interaction experience (Social retrieval). (**B**) Time course of freezing during CFC1 and CFC2. (**C**) Mice in the Fear retrieval group freeze significantly more during CFC2 than during CFC1. (**D**) Time course of mouse and cup investigation time during SI1. (**E**) Time course of mouse and cup investigation time during SI2. (**F**) In SI1, mice spend more time investigating the female mouse compared to the cup. In SI2, there is no difference in the time investigating the female mouse and the cup. (**G**) Mice traveled a shorter distance during SI2 when compared to SI1. (**H**) Representative image of EYFP^+^, mKate2^+^ and c-fos^+^ ensembles in the dDG. The white box delineates the area magnified in the right panels. The white arrows point to a mKate2^+^ c-fos^+^ cell. (**I**) The percentage of EYFP^+^ and mKate2^+^ cells active during exposure to shock and during exposure to a female. (**J**) The percentage of c-fos^+^ cells does not differ between the Fear retrieval and Social retrieval group. (**K**) Diagram showing between-group comparisons. (**L**) The percentage of EYFP^+^ cells that also express mKate2 does not differ between the Fear retrieval and Social retrieval group. (**M**) The Fear retrieval group has a higher percentage of EYFP^+^ cells that also express c-fos when compared to the Social retrieval group. (**N**) The Social retrieval group has a higher percentage of mKate2^+^ cells that also express c-fos when compared to the Fear retrieval group. (**O**) Diagram showing within-group comparisons. (**P**) The overlap between ensembles active 1 d apart is greater than that of ensembles active 6 or 5 d apart for both Fear retrieval and Social retrieval groups. The overlap between ensembles 1 d apart for the Fear retrieval group is higher than that of the Social retrieval group. n = 6-7 mice per group. Error bars show mean ± SEM. *p < 0.05, **p < 0.01, ****p < 0.0001. dDG, dorsal dentate gyrus; EYFP, enhanced yellow fluorescent protein; 4-OHT, 4-hydroxytamoxifen; DOX, doxycycline; CFC, contextual fear conditioning; SI, social interaction; Sac, sacrifice.

To assess how ensembles are reactivated across time, we compared the normalized ensemble reactivation rates in a within-group manner **(Figure 6O)**. Surprisingly, for both groups, the reactivation of the ensembles active 1 day apart was greater than that of the ensembles 6 or 5 days apart, regardless of whether the experiences 1 day apart were of similar or distinct valence **(Figure 6P).** We observed that the reactivation of ensembles active 1 day apart was greater for the Social retrieval group, which underwent two social experiences, than that of the Fear retrieval group, which underwent a social experience and CFC. These results indicate that the ensembles of valenced experiences are modulated by time.

## DISCUSSION

Here, we generated a novel IEG-tagging system, the mArc system, to tag multiple *Arc*^+^ ensembles in a single mouse and examine how various memories are encoded in the brain. This system allows for the labeling of two previously activated *Arc* ensembles and a third activated ensemble visualized through immunohistochemistry. Using the mArc system, we investigated how multiple contextual experiences are encoded in neuronal ensembles in the DG, a region important for contextual memory. We found that similar contextual and valenced experiences are encoded in overlapping DG ensembles. Moreover, we uncovered a significant role of time in ensemble reactivation, where experiences close in time are stored in more similar ensembles compared to those further apart in time. Our data highlight the dynamic nature of DG ensembles and suggest that DG ensembles bind contextual and temporal information to represent individual experiences.

Initial attempts were focused on developing a dual-tagging system that allowed for *Arc* and *c-fos* ensembles to be tagged in a within-mouse design. However, the low level of overlap between the tagged *Arc* and *c-fos* ensembles, along with recent work showing distinct roles for IEG-specific ensembles^18^, led us to develop an *Arc*-based tagging system. This approach is a powerful tool to investigate the unique properties of multiple *Arc* ensembles and how these ensembles are reactivated without the confound of different IEG pathways. Nevertheless, our *Arc* and *c-fos* experiments add to the growing literature regarding differences in IEG expression and highlight the need to investigate the functions of distinct IEG ensembles. Although the c-fos-tTA strategy in the quadruple transgenic mouse underestimated the number of cells in the DG **(Figure S1)**, this strategy could still be useful for comparing *Arc* and *c-fos* ensembles in other brain regions. Recently, a *c-fos*-based dual-tagging system was generated to investigate how multiple valenced experiences are encoded in ventral CA1^35^. While this system combined tagging strategies based on *c-fos*^16,36^, our system focused on *Arc* due to its significant role in long-term memory and in various forms of synaptic plasticity^20,37^. Overall, these dual-tagging systems provide the field with powerful genetic tools to access multiple active ensembles in a single mouse.

Although both the *Arc* tagging strategies that make up the mArc system label a relatively similar percentage of cells in the DG during contextual experiences (∼2-6%), there are slight differences between the two components. While the ArcCreER^T2^ strategy labels cells indelibly through Cre-mediated recombination, the *Arc*-RAM strategy labels cells persistently for 2 weeks before the reporter fluorescence starts degrading^14^. Despite these differences, the mArc system provides enduring labeling of neural ensembles. Moreover, the *Arc*-RAM strategy consistently labels a larger percentage of cells than the ArcCreER^T2^ strategy. This labeling difference could be attributed to: 1) differences in the length of the tagging windows (i.e., a single injection of 4-OHT versus removal of DOX for 3 days); 2) a transgenic versus viral strategy; or 3) varied promoter lengths (i.e., a full *Arc* promoter versus a short, synthetic promoter). Regardless of these differences, we can successfully label ensembles that have similar synaptic properties, providing evidence of a tagging system that labels multiple behaviorally relevant *Arc* ensembles in the same mouse.

An advantage of the mArc strategy is that it allows for the identification and access to multiple *Arc* ensembles for further physiological characterization. Our results characterizing the synaptic and cellular properties of *Arc* ensembles show that these ensembles receive a larger number of excitatory input compared to unlabeled cells, which are in line with previous studies demonstrating that *Arc^+^* and *c-fos^+^*ensembles exhibit increased synaptic connectivity after learning^18,38–41^. While these studies recorded from cells hours to days after learning, we recorded from cells 5-7 days after CFC, indicating that this synaptic signature persists significantly after learning in a timeline comparable to that of our behavioral exposures. Whether this increased synaptic connectivity happens selectively between *Arc* ensembles across brain regions remains to be determined^42^. In addition, we found that labeled *Arc* ensembles do not exhibit differences in measures of intrinsic excitability when compared to unlabeled neurons. Previous work has shown that neuronal excitability is increased transiently (in the scale of hours to 2 days) after memory encoding or retrieval^41,43–46^, and these results are consistent with our findings since we record from cells past the excitability window reported in these studies.

In the DG, ensembles are reactivated in a context-specific manner^4,14,47,48^, and our work expands on previous research by demonstrating that this context-specific reactivation persists across multiple contextual exposures. Surprisingly, across a variety of experimental paradigms, we observed a strong time-dependent effect on the reactivation of ensembles, where experiences closer in time were encoded in more overlapping ensembles. Notably, when contextual experiences are similar, we always observed this effect. However, when comparing ensembles across different contextual experiences, this time-dependent effect is contingent upon the *valence* of experiences. For distinct neutral experiences, we do not observe a time-dependent modulation of ensembles. For distinct valenced experiences, we observed that the ensembles active 1 day apart had a higher reactivation rate than those 5 or 6 days apart, even when the experiences 1 day apart were of opposite valence. A possible explanation for these results could be memory linking, where salient memories acquired close in time become associated through overlapping ensembles^43,49^. It is worth noting that the reactivation of valenced ensembles might not be driven by valence alone, since similar contextual cues are present across experiences of the same valence. Further work is needed to investigate how interactions between contextual and valence information modulate hippocampal ensembles.

The effect of time observed in our studies is similar to representational drift, the phenomenon where neuronal representations change over time despite no changes in the environment or behavior^50,51^. *In vivo* recordings of the hippocampus have shown that population representations of an environment gradually change over time^52–56^. While the degree to which representations change through time varies across hippocampal subregions, studies show that DG granule cell stably encode contextual information across days^54,57^. In contrast, our results show that only a small fraction of contextual ensembles (2-8%) is reactivated during subsequent exposures and that ensembles are strongly modulated by time. Future work that examines the relationship between neuronal firing rates/calcium activity and IEG expression^58–60^ will be crucial for bridging the discrepancies between engram and *in vivo* recording studies.

Aside from being modulated by context, DG ensemble reactivation has been associated with memory strength, as seen in the case of fear extinction^3,7,61^, pharmacological interventions^62^, and disease states^63,64^. Curiously, we did not see any differences in ensemble reactivations between mice that underwent 1 or 2-Trial CFC despite observing changes in the relative strength of the memory. A potential reason for the lack of differences between these groups could be that the strength of the memory is still relatively high for both groups. Another explanation could be that DG ensembles do not encode the strength or valence of the memory *per se*, but form representations of contextual and temporal information that then gets integrated with strength or valence information in downstream brain regions, such as the amygdala^65,66^.

In summary, the mArc strategy has allowed us to investigate how multiple experiences are encoded in the hippocampus and to study the impact of context, valence, and time on ensemble reactivation in a within-mouse design. These studies have allowed us to investigate the evolution of engrams over time and begin to determine the impact of neuronal drift on ensemble activity. This new genetic tool will be helpful in characterizing engrams as representations are formed and evolve over time not only in normal memory storage, but also in disease states where memories either degrade, such as in Alzheimer’s disease (AD) and age-related cognitive decline (ARCD), or become maladaptive, such as in post-traumatic stress disorder (PTSD) and generalized anxiety disorder (GAD).

## ACKNOWLEDGMENTS

We thank Dr. Taiga Abe, Dr. Michael Drew, and Michelle Jin for comments on the manuscript, Dr. Steve Ramirez for feedback on the manuscript and for providing viral constructs, Dr. Clay Lacefield for comments on the manuscript and for aiding with technical expertise, as well as Drs. Gergely Turi and Victor Luna for generously providing technical and statistical expertise. We thank Drs. Steve Siegelbaum, René Hen, and Dani Dumitriu for insightful comments on the project, and Drs. Alcino Silva and Johannes Gräff for providing transgenic mouse lines. M.S. was supported by NINDS 1F99NS129178-01, NIMH 1F31MH125656-01A1, and NICHD 5T32HD007430-20. N.E.B. was supported by NSF GRFP DGE-2036197. Y.L. was supported by R01NS115543 and R01MH050479. C.A.D. was supported by a Whitehall Foundation Grant, an NIH Transformative Award 1R01HD101402-01, and DP5OD017908.

## AUTHOR CONTRIBUTIONS

Conceptualization, M.S. and C.A.D.; Investigation, M.S., T.D.Y., M.M., J.N., N.E.B., N.V., E.Z.; Resources, X.S., Y.L.; Writing – Original Draft, M.S., C.A.D.; Writing – Review & Editing, M.S., C.A.D.; Supervision, C.A.D; Funding Acquisition, C.A.D.

## INCLUSION AND DIVERSITY

We support inclusive, diverse, and equitable conduct of research.

## RESOURCE AVAILABILITY

### Lead contact

Further information and request for resources and reagents should be directed to the lead contact, Christine Ann Denny.

### Materials availability

Plasmids generated in this study will be deposited to Addgene.

### Data and code availability

- Microscopy and all other data reported in this paper will be shared by the lead contact upon request.
- This paper does not report original code.
- All data and any additional information required to reanalyze the data reported in this paper is available from the lead contact upon request.

## EXPERIMENTAL MODEL AND STUDY PARTICIPANT DETAILS

### Mice

ArcCreER^T2^ (+) x R26R-STOP-floxed-EYFP^67^ homozygous female mice were bred with R26R-STOP-floxed-EYFP homozygous male mice to generate experimental ArcCreER^T2^ (+) x R26R-STOP-floxed-EYFP mice. For electrophysiology experiments, ArcCreER^T2^ (+) mice were used. ArcCreER^T2^ mice have been backcrossed for more than 10 generations onto a 129S6/SvEv line.

ArcCreER^T2^ (+) x R26R-STOP-floxed-EYFP homozygous x c-fos-tTA mice were generated by breeding ArcCreER^T2^ (+) x R26R-STOP-floxed-EYFP homozygous mice with c-fos-tTA mice. c-fos-tTA mice were obtained from Dr. Alcino Silva’s laboratory and were initially on a C57BL/6N background. These mice were backcrossed onto a 129S6/SvEv line for at least 5 generations before breeding with the ArcCreER^T2^ mice.

Quadruple transgenic mice were generated by breeding ArcCreER^T2^ (+) x R26R-STOP-floxed-EYFP homozygous x c-fos-tTA mice with tetO-tdTomato mice and were raised on a diet containing 40 mg/kg DOX. tetO-tdTomato mice were obtained from Dr Johannes Gräff’s laboratory and were initially on a C57BL/6N background. These mice were backcrossed onto a 129S6/SvEv line for at least 5 generations.

Ovariectomized female 129S6/SvEv mice were purchased from Taconic at 5-7 weeks of age.

All mice were housed 2-5 per cage in a 12 h (6AM to 6PM) light/dark cycle colony room at 22°C. Food and water were provided *ad libitum*. Experiments were performed during the light phase. Mice were 2-5 months of age at the start of the experiment. All experimental procedures were approved by the Institutional Animal Care and Use Committee (IACUC) at the New York State Psychiatric Institute and Columbia University.

## METHOD DETAILS

### Viruses

AAV9-*Arc*-RAM-mKate2 (AAV9-ESARE-d2tTA-TRE-mKate2) was produced in the laboratory of Dr. Yingxi Lin. The AAV was generated in HEK293T cells and purified using an adapted gradient purification protocol as previously described^14^. The AAV vector was serotyped with AAV9 coat proteins and packaged at Vigene Biosciences. For most experiments, AAV9-*Arc*-RAM-mKate2 (titer: 5.11 x 10^13^ GC/mL) was diluted 1:19 in saline.

The pAAV-TRE-mCherry plasmid was constructed as previously described^47^ and packaged at the University of Massachusetts Medical School Gene Therapy Center and Vector Core to generate AAV9-TRE-mCherry virus. For *Arc* and *c-fos* ensemble tagging, AAV9-TRE-mCherry (titer: 7.9 x 10^12^ GC/mL) was diluted 3:1 in saline.

For electrophysiology experiments where two *Arc* ensembles were labeled simultaneously, AAV9-*Arc*-RAM-mKate2 and AAV5-Ef1a-DIO-EYFP-WPRE-pA were diluted 1:12:7 in saline and subsequently co-injected. AAV5-Ef1a-DIO-EYFP-WPRE-pA was purchased from UNC Vector Core.

### Drugs

#### 4-hydroxytamoxifen (4-OHT)

Recombination was induced by 4-OHT as previously described^4,68^. 4-OHT was dissolved by sonication in 10% ethyl alcohol/90% corn oil at a concentration of 10 mg/mL. Mice were administered an injection of 0.2 mL intraperitoneally (i.p.) 5 h prior to behavioral testing. After an injection, mice were dark housed for 3 days in order to limit additional stimuli that might induce *Arc* expression^4^.

#### Kainic acid (KA)

For seizure induction, KA was dissolved in saline at a concentration of 2 mg/mL and injected i.p. at 20 mg/kg^69^. Mice were injected with KA and monitored continuously for behavioral onset of seizure for at least 3 hours post-injection and daily for the next 3 days. Only mice which exhibited full motor seizures were considered for analysis.

#### Doxycycline (DOX)

To prevent tTA-mediated expression, DOX was given at 40 mg/kg in the mouse diet^6^.

### Stereotaxic surgeries

Mice were anesthetized with isoflurane (4% induction, 1-3% maintenance in O_2_) and head-fixed in a stereotactic frame (David Kopf Instruments). Ophthalmic ointment was used to lubricate the eyes and a heating pad was used to maintain the body temperature at 37°C. Carprofen was injected subcutaneously prior to surgical procedures, immediately afterward, and daily for at least 3 d following surgery. The surgical area was shaved and disinfected prior to incision.

Viruses were infused unilaterally (50 nL per infusion, 30 s between infusions) into the DG or CA3 using a glass pipette connected to a Nanoject II injector. After each infusion, the glass pipette was kept in place for 5 min before removal to allow viral diffusion. The coordinates relative to Bregma and viral volumes were: DG (AP –1.50 mm, ML ±1.05 mm, DV –1.95 mm; 200 nL per site) and CA3 (AP –1.46 mm, ML ±2.00 mm, DV –2.10 mm; 250 nL per site). Mice injected with AAV9-*Arc*-RAM-mKate2 were put on DOX diet 1 d before or the day of surgery, while mice injected with AAV9-TRE-mCherry were put on DOX diet at least 5 d before surgery. For DG and CA3 injections, only mice with a viral infection of more than 80% or 70% of the area of interest, respectively, were included in the analysis.

### Behavior

#### Context exposure

Mice were transported into the behavior room and placed in context A, which consisted of a 7 x 7 x 12 in chamber with clear plastic front and back walls, stainless steel walls on each side, and stainless-steel bars on the floor. Each chamber was located inside a larger, insulated plastic cabinet that provided protection from outside light and noise. The context was scented with lemon scent. Mice were allowed to explore the chamber for 3 min and their behavior was recorded. The chambers were cleaned with 70% ethanol between each run.

Mice exposed to context B were transported to the behavior room inside a white container. Context B was a modified context A, where the walls of the chamber were covered with round, colored plastic inserts to change the shape of the chamber. The context was scented with anise scent. The room lights were off, and a red light was used for illumination. The chambers were cleaned with Virkon-S solution between each run.

Mice exposed to context C were transported in their home cage to a different room than context A and B. Context C consisted of a chamber with the same dimensions to context A, but the floor was covered with a plastic insert, bedding was placed on top, and colored plastic inserts were used to create semi-circular walls inside the chamber. No scent was used in context C. The chambers were cleaned with Sani-Cloth disposable wipes between each run. The different features used for context A, B, and C are listed in **Table S1**.

#### Contextual Fear Conditioning (CFC)

A 3-shock CFC paradigm was administered as previously described^4,63,70^. Mice were placed in the conditioning chamber and received three shocks 180, 240, and 300 s later (2 s, 0.75 mA). Mice were removed from the chamber 15 s following the last shock. During re-exposure, mice were placed in the conditioning chamber for 3 minutes and did not receive any shocks. Freezing was scored using FreezeFrame4.

#### Social Interaction (SI)

The SI assay was modified from a previously published protocol^63^. Mice were individually housed overnight prior to testing. Mice were placed in a large open field arena containing two upside-down wire mesh cylinder cups on opposite ends. The testing arena was a white, plastic transport box (55 x 40 x 15 cm). During this 10 min exposure, one cup remained empty while the other cup held a previously unencountered OVX female mouse. After testing, male mice and their respective OVX female mice were co-housed for the remainder of the experimental protocol.

For experiments where two SI tests occurred, the second SI test consisted of separating the co-housed male and female mice for 1 h. Then, male mice were re-exposed to the testing arena as described above. During this 10 min exposure, one cup remained empty while the other cup held the now familiar OVX female mouse. The time spent exploring each cup and the total distance traveled were automatically quantified using ANY-maze software. Mice that failed to explore both cups during a social interaction assay were excluded from the analysis.

### Ensemble labeling

For double-labeling *Arc*^+^ experiments, mice were placed into a dark, separate housing room in a fresh cage the night before 4-OHT injection. The next day, to open the labeling window for the ArcCreER^T2^ strategy, mice were injected with 4-OHT intraperitoneally 5 h before behavior. Following behavior, mice were housed in a dark, separate housing room for 3 days. After, mice were taken out of the dark room, placed in a clean cage, and returned to the colony room.

To open the labeling window for the *Arc*-RAM strategy, DOX diet was removed from mice and replaced with normal chow 48 h before behavioral testing. Mice were returned to a DOX diet 24 h after behavioral testing, if needed.

For simultaneous *Arc*^+^ and *c-fos*^+^ labeling experiments, mice followed the protocol described above for tagging except they were turned to a DOX diet immediately after behavioral testing.

For experiments where only *Arc*-RAM labeling was performed, DOX diet was removed from mice and replaced with normal chow 48 h before behavioral testing. Mice were returned to a DOX diet 24 h after behavioral testing, if needed.

### Immunohistochemistry

Mice were deeply anesthetized with (*R*,*S*)-ketamine/xylazine (100 mg/kg and 10 mg/kg, respectively) and transcardially perfused with 1X phosphate buffered saline (PBS), followed by 4% paraformaldehyde (PFA) in 1X PBS. Brains were extracted and post-fixed overnight in 4% PFA at 4°C and then transferred to 1X PBS at 4°C until further use. Coronal sections of the hippocampus were cut (50 μm) using a vibratome and stored in 1X PBS with 0.1% NaN_3_ until immunohistochemistry was performed. Free-floating sections were washed 3 times, 10 min each, in 1X PBS and blocked in 10% normal donkey serum (NDS) in 1X PBS and 0.5% Triton X-100 (PBST) for 2 h at room temperature. Sections were then incubated with primary antibody in PBST overnight at 4°C. The next day, sections were washed 3 times, 10 min each, in 1X PBS and incubated in secondary antibody in PBS for 2 h at room temperature. Then, sections were washed 3 times, 10 min each, in 1X PBS. Some sections were further incubated in Hoechst 33342 (1:1,000) in 1X PBS for 10 min and washed again 3 times, 10 min each, in 1X PBS. Sections were mounted onto glass slides and coverslipped with Fluoromount-G. Primary and secondary antibodies and their corresponding concentrations are listed in **Table S2.**

### Confocal microscopy and cell counting

For quantification of fluorescently labeled cells, every sixth section along the longitudinal axis of the hippocampus was imaged and cells in the granule cell layer of the DG or pyramidal layer of CA3 were counted exhaustively for each mouse (4-7 sections per mouse). Z-stacks of immunolabeled sections were obtained using a confocal scanning microscope (Leica TCS SP8) equipped with two simultaneous PMT detectors. Fluorescence from Cy2 was excited at 488 nm and detected at 500-550 nm, fluorescence from Cy3 was excited at 552 nm and detected at 588-638 nm, and fluorescence from Alexa Fluor 647 was excited at 634 nm and detected at 650-700 nm. Sections were imaged with a dry Leica 20x objective (numerical aperture 0.70, working distance 0.5 mm), a pixel size of 1.08 x 1.08 μm, a z-step of 3 μm, and a z-stack of 12 μm. Several fields of view were stitched together to form tiled images by using an automated stage as well as the tiling function and algorithm of the LAS X software.

To obtain a Hoechst^+^ density estimate, 4 z-stacks, 12 μm thick each, were obtained with a 63x objective at different locations of the dentate gyrus for a total of 4 mice. The number of Hoechst^+^ cells in each z-stack was quantified and divided by the region’s volume (area x 12 μm). The Hoechst^+^ cells/volume measurements were averaged across sections and mice to provide a final estimate of Hoechst density in the DG. The same process was repeated in CA3 to obtain a Hoechst density estimate for the region. This method was also used to quantify the overlap between EYFP^+^ and mKate2^+^ cells labeled during a kainic-acid induced seizure.

Fluorescently-labeled cells in the granule cell layer of the DG or pyramidal layer of CA3 were manually counted for each acquired z-stack using ImageJ by an investigator blind to treatment. To identify double or triple-labeled cells, we first identified and quantified fluorescently-labeled cells one channel at a time and then checked that the cells identified in multiple channels were double– or triple-labeled. To calculate overlap/chance measurements, we divided the proportion of observed co-labeled cells (co-labeled^+^/Hoechst^+^) by the expected chance overlap between labeled ensembles (Label 1^+^/Hoechst^+^) x (Label 2^+^/Hoechst^+^).

### Whole-cell patch-clamp electrophysiology

#### Slice preparation

Electrophysiology was performed as previously described^18^. Eight to nine-week-old mice were used for electrophysiology studies. Neural ensembles were tagged as forementioned following a 3-shock CFC paradigm. Five to seven days later, mice were sacrificed and coronal slices (300 µm) of the dorsal hippocampus were prepared using in carbogenated (95% O_2_/5% CO_2_) ice-cold cutting solution containing (in mM) 210 sucrose, 2.5 KCl, 1.2 NAH_2_PO_4_, 10 MgCl_2_, 0.5 CaCl_2_, 26 NaHCO_3_, 25 glucose, 20 HEPES, 5 sodium-L-ascorbate, 2 thiourea, and 3 Na pyruvate, ∼340 mOsm osmolarity, pH 7.3 using a vibratome (VT1200, Leica). Slices were transferred to a carbogenated recovery solution at 32°C for 15 min. The solution consisted of 50% cutting solution and 50% artificial cerebrospinal fluid (ACSF), which was composed of (in mM) 119 NaCl, 2.5 KCl, 1.2 NaH_2_PO_4_, 2 MgCl_2_, 2 CaCl_2_, 26 NaHCO_3_, and 10 glucose, 305 mOsm osmolarity, pH 7.3. After, slices were transferred to a chamber containing room temperature (∼23°C) ACSF and kept in ACSF for at least 45 min before recording.

#### Whole-cell recordings

Recordings were performed in a chamber perfused with carbogenated ACSF at room temperature at a flow rate of 2 mL/min. Borosilicate glass pipettes with a tip resistance of 3-5 MΩ were used. mEPSC recordings were obtained from cells voltage-clamped at –70 mV in ACSF containing 0.5 µM tetrodotoxin (TTX, Tocris) and 50 µM picrotoxin (PTX, Tocris). A Cs-based internal solution was used containing (in mM) 130 CsMeSO3, 10 phosphocreatine, 1 MgCl2, 10 HEPES, 0.2 EGTA, 4 Mg-ATP, 0.5 Na-GTP, 295 mOsm osmolarity, pH adjusted to 7.25 with CsOH. Passive membrane properties were recorded with the following internal solution (in mM): 130 K^+^-D-gluconate, 10 KCl, 10 HEPES, 4 Mg-ATP, 0.5 Na-GTP, 0.2 EGTA, 1 MgCl2, pH adjusted to 7.25 with CsOH, 295 mOsm osmolarity with ACSF containing 50 µM PTX, 50 µM APV and 20 µM DNQX.

Slices were visualized using an Olympus BX51WI microscope with a 40x objective lens. Recordings were restricted to neurons in the granule cell layer of the dorsal dentate gyrus. Neurons expressing EYFP, mKate2, or both were visualized with a PRIOR Lumen 200 LED light and selected for targeted patch-clamp recordings.

#### Data acquisition and analysis

Data were acquired with a Multiclamp 700B (Molecular Devices), filtered at 3 kHz and digitized at 10 kHz with Digidata 1440A and Clampex 10.2 software (Molecular Devices). Recordings were discarded if access resistance was greater than 30 MΩ or changed more than 15% throughout the recordings. Data were analyzed using MiniAnalysis (Synaptosoft) and Clampfit 10.2 (Molecular Devices).

## QUANTIFICATION AND STATISTICAL ANALYSIS

All data were analyzed using Prism 10. Means of two groups were compared using two-sided paired or unpaired Student’s *t*-test. One-way ANOVA, repeated measures (RM) one-way ANOVA, or the Kruskal-Wallis test were used to compare the means of three or more groups, when appropriate. Two-way RM ANOVA was used to compare group means when there were two or more variables. Significant effects were followed by Tukey’s or Dunn’s multiple comparisons test. RM ANOVAs were calculated with Geisser-Greenhouse correction. Wilcoxon and Friedman tests were used when appropriate. Alpha was set to 0.05 for all analyses. All data are presented as mean ± SEM. The statistical details of all experiments can be found in **Table S3**.

## ADDITIONAL RESOURCES

**Figure design and visualization**

Figures and graphical illustrations were created with Biorender (Biorender.com).

## SUPPLEMENTAL TEXT AND FIGURES

**Figure S1.**
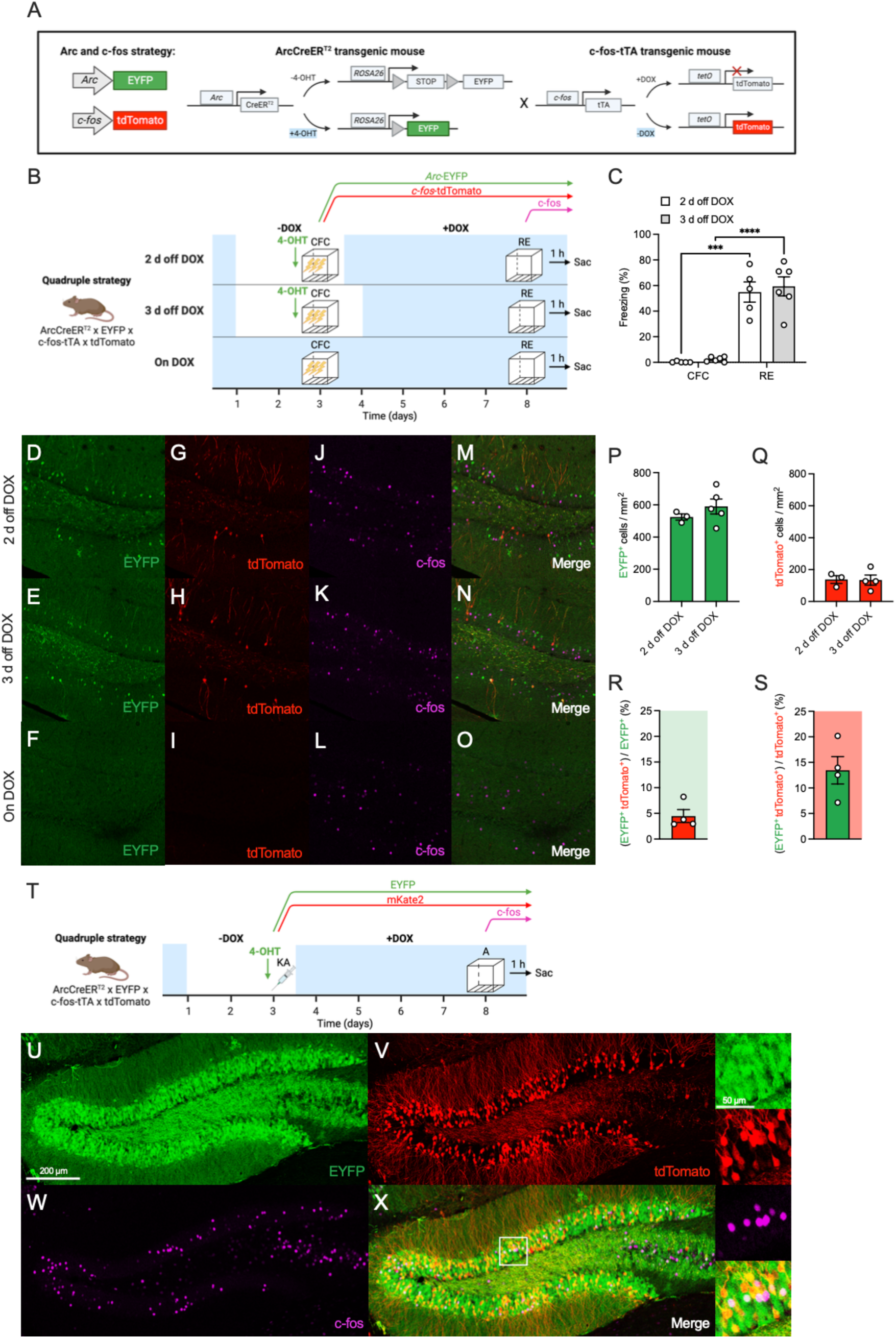
Insufficient labeling of an active *c-fos*^+^ ensemble using an Arc and c-fos quadruple strategy, related to Figure 1. (**A**) Genetic strategy to label an *Arc* and a *c-fos* ensemble in the same mouse by breeding the transgenic ArcCreER^T^^2^ x EYFP and c-fos-tTA x tdTomato mouse lines together. (**B**) Experimental design to label an *Arc* and a *c-fos* fear ensemble in the DG. (**C**) Mice administered a 3-shock CFC paradigm do not freeze during training (i.e., before shocks are delivered). Mice freeze significantly during re-exposure and there is no difference in the freezing levels during re-exposure for the mice taken off DOX for 2 or 3 days. Representative image showing: (**D-F**) EYFP^+^ cells, (G-I) tdTomato^+^ cells, (J-L) c-fos^+^ cells, and (**M-O**) merged DG section. (**P**) The number of EYFP^+^ cells is comparable between the groups taken off DOX for 2 or 3 days. (**Q**) The number of tdTomato^+^ cells is comparable between the groups taken off DOX for 2 or 3 days. (**R**) The percentage of EYFP^+^ cells that also express tdTomato for the group taken off DOX for 3 days. (**S**) The percentage of tdTomato^+^ cells that also express EYFP for the group taken off DOX for 3 days. (**T**) Experimental design to label *Arc* and *c-fos* ensembles activated after a kainic acid injection. Representative image showing: (**U**) EYFP^+^ cells, (**V**) tdTomato^+^ cells, (**W**) c-fos^+^ cells, and (**X**) merged DG section. n = 3-4 mice per group. Error bars show mean ± SEM. ***p < 0.001, ****p < 0.0001. EYFP, enhanced yellow fluorescent protein; 4-OHT, 4-hydroxytamoxifen; tTA, tetracycline-controlled transactivator; DOX, doxycycline; tetO, tetracycline operator; CFC, contextual fear conditioning; RE, re-exposure; Sac, sacrifice; KA, kainic acid.

**Figure S2.**
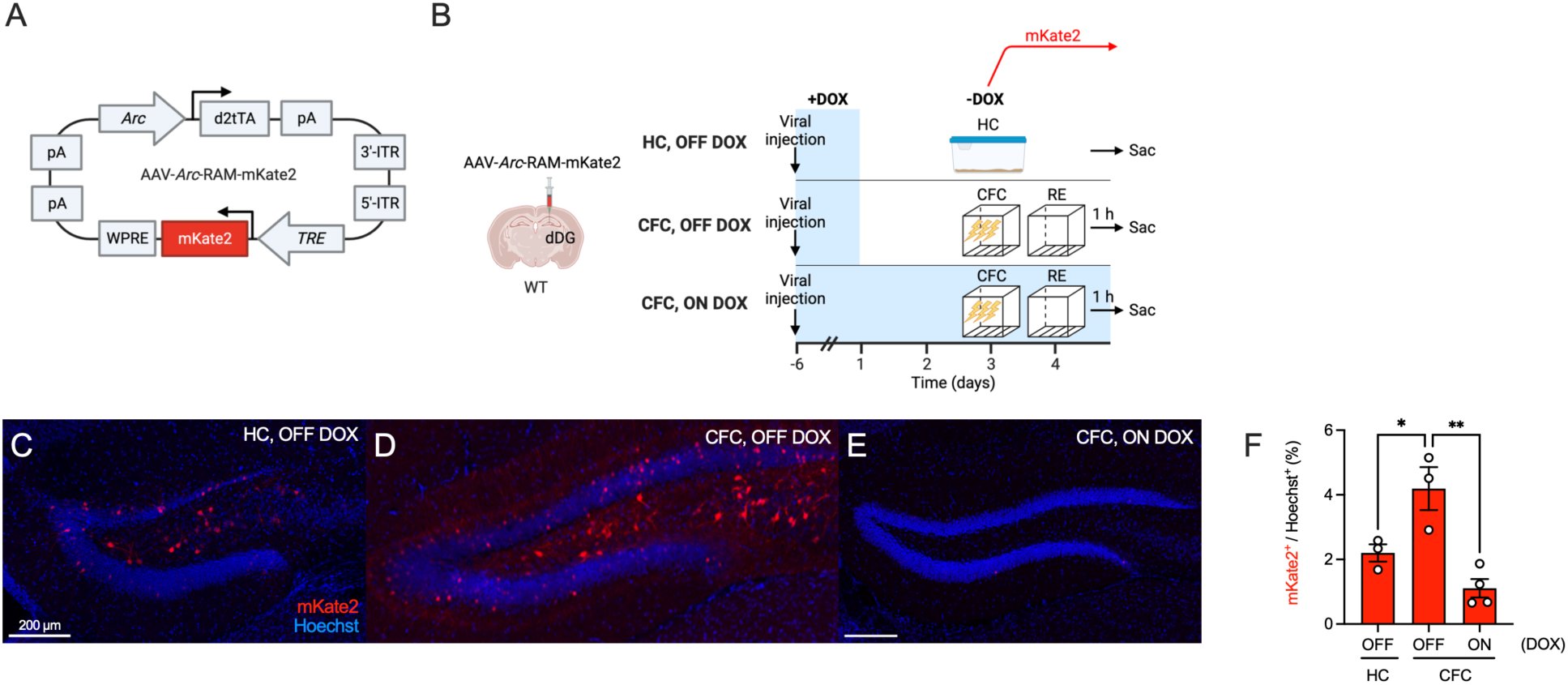
Labeling of an *Arc*^+^ ensemble in the dentate gyrus using a novel *Arc*-RAM strategy, related to Figure 1. (**A**) Genetic design of the *Arc*-RAM vector. (**B**) Experimental design to label active ensembles using the *Arc*-RAM strategy. Representative images showing: (**C**) mKate2^+^ cells for the OFF DOX, HC group, (**D**) mKate2^+^ cells for the OFF DOX, CFC group, (**E**) mKate2^+^ cells for the ON DOX, CFC group. (**F**) The percentage of mKate2^+^ cells in the OFF DOX, CFC group is greater than in the OFF DOX, HC or ON DOX, CFC groups. n = 3-4 mice per group. Error bars show mean ± SEM. *p < 0.05, **p < 0.01. d2tTA, destabilized tetracycline transactivator; TRE, tetracycline response element; dDG, dorsal dentate gyrus, WT, wild type; DOX, doxycycline; CFC, contextual fear conditioning; RE, re-exposure; HC, home cage.

**Figure S3.**
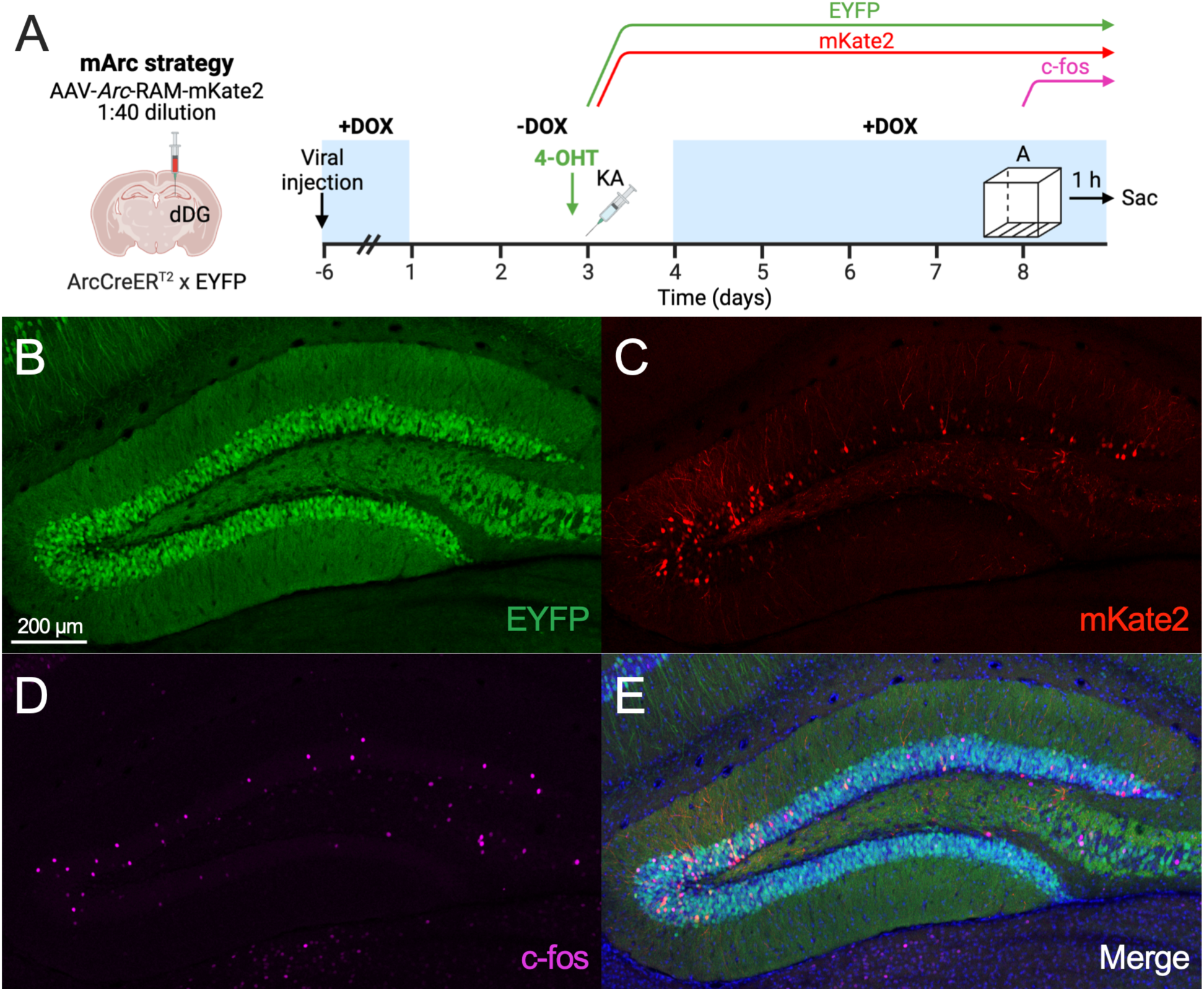
Dense labeling of *Arc* ensembles following a kainic acid-induced seizure in the ArcCreER^T2^ x EYFP strategy, but not in the Arc-RAM viral strategy when injected at a lower titer, related to Figure 2. (**A**) Experimental design to label an *Arc* ensemble activated by kainic acid using a lower viral titer of *Arc*-RAM. Representative images showing: (**B**) EYFP^+^ cells, (**C**) mKate2^+^ cells, (**D**) c-fos^+^ cells, and (**E**) merged DG section. EYFP, enhanced yellow fluorescent protein; dDG, dorsal dentate gyrus; DOX, doxycycline; 4-OHT, 4-hydroxytamoxifen; KA, kainic acid; Sac, sacrifice.

**Figure S4.**
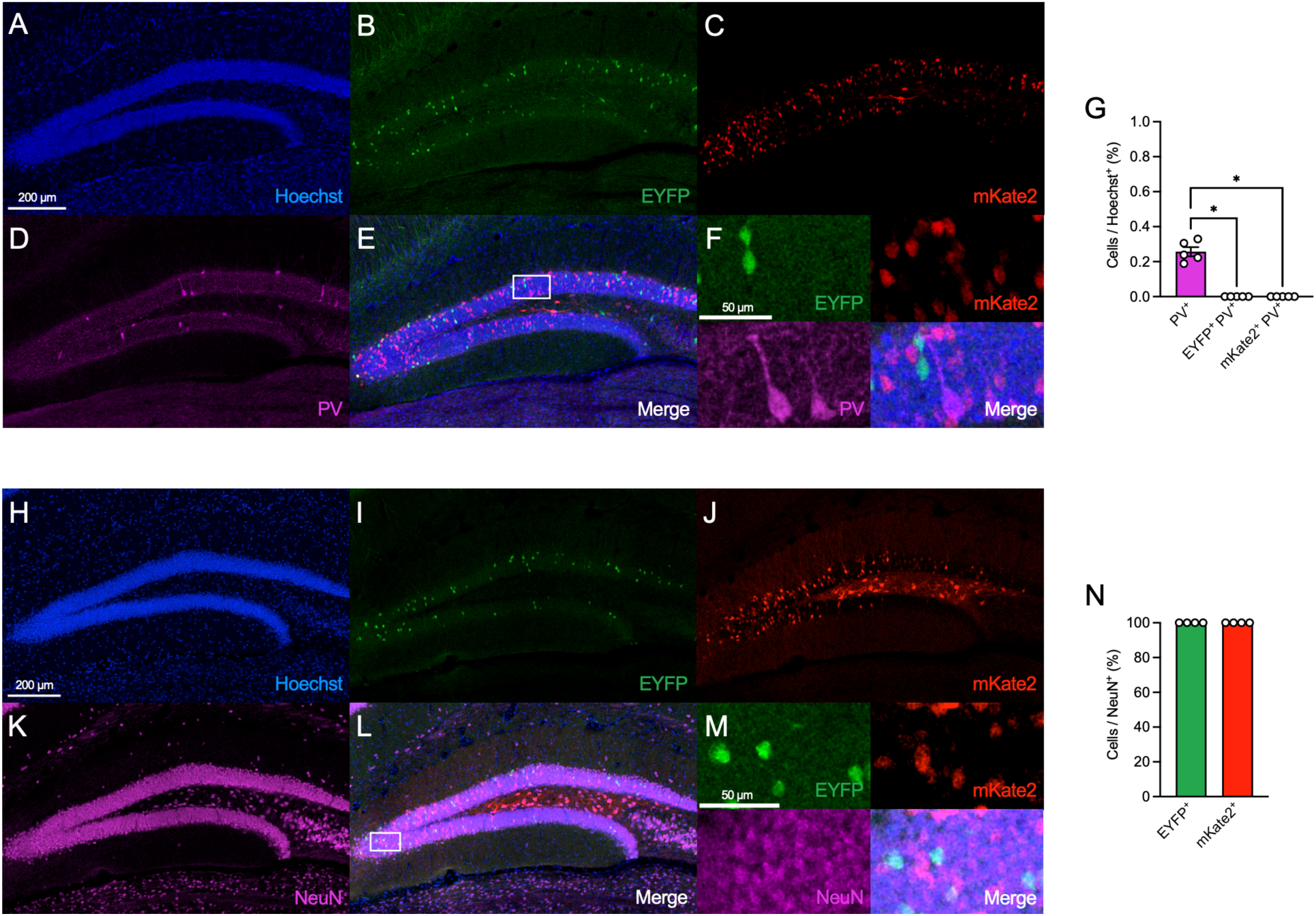
Characterization of ensembles labeled with the mArc strategy, related to Figure 3. Representative images showing labeling of: (**A**) Hoechst^+^ cells, (**B**) EYFP^+^ cells, (**C**) mKate2^+^ cells, (**D**) PV^+^ cells, and (**E**) merged DG section. The white box shows the area magnified in F. (**F**) Magnified images showing EYFP^+^, mKate2^+^, and PV^+^ cells. (**G**) The percentage of cells quantified in the DG. The EYFP^+^ and mKate2^+^ ensembles do not overlap with a PV^+^ ensemble. Representative images showing labeling of: (**H**) Hoechst^+^ cells, (**I**) EYFP^+^ cells, (**J**) mKate2^+^ cells, (**K**) NeuN^+^ cells, and (**L**) merged DG section. The white box shows the area magnified in M. (**M**) Magnified images showing EYFP^+^, mKate2^+^, and NeuN^+^ cells. (**N**) The EYFP^+^ and mKate2^+^ ensembles completely overlap with NeuN. n = 4-5 mice. Error bars show mean ± SEM. *p < 0.05. EYFP, enhanced yellow fluorescent protein; PV, parvalbumin; NeuN, neuronal nuclei.

**Figure S5.**
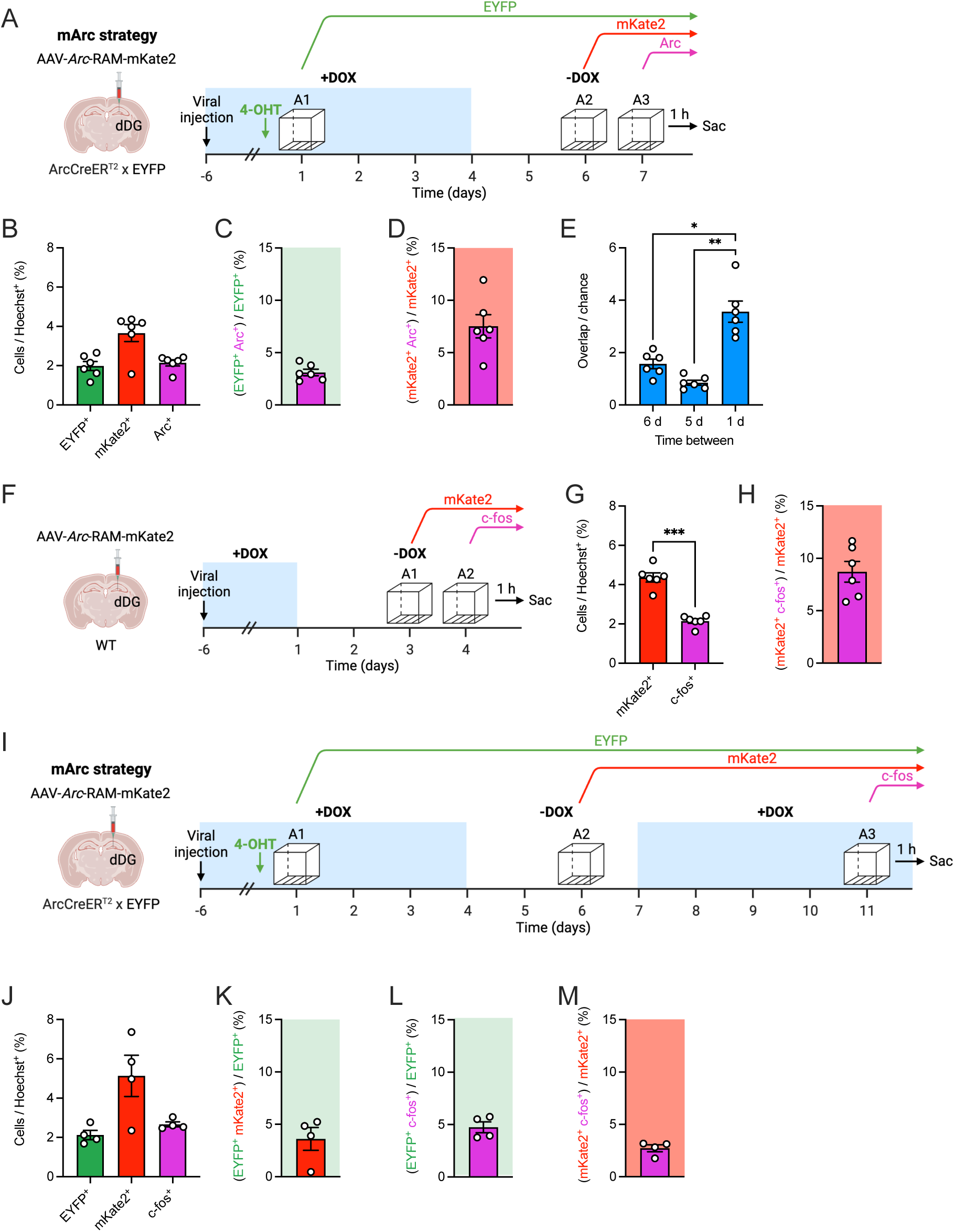
Ensemble reactivation is modulated by time between experiences, related to Figure 4. (**A**) Experimental design to label the ensembles active during three exposures to the same context. (**B**) The percentage of EYFP^+^, mKate2^+^ and Arc^+^ cells. (**C**) The percentage of EYFP^+^ cells that also express Arc. (**D**) The percentage of mKate2^+^ cells that also express Arc. (**E**) The overlap between ensembles active 1 d apart is greater than that of ensembles active 6 d or 5 d apart for the Same Context group. (**F**) Experimental design to label the ensembles active during two exposures to the same context, with one day in between exposures. (**G**) The percentage of mKate2^+^ and c-fos^+^ cells. (**H**) The percentage of mKate2^+^ cells that also express c-fos. (**I**) Experimental design to label the ensembles active during three exposures to the same context, with five days in between exposures. (**J**) The percentage of EYFP^+^, mKate2^+^ and c-fos^+^ cells. (**K**) The percentage of EYFP^+^ cells that also express mKate2. (**L**) The percentage of EYFP^+^ cells that also express c-fos. (**M**) The percentage of mKate2^+^ cells that also express c-fos. n = 4-6 mice per group. Error bars show mean ± SEM. *p < 0.05, **p < 0.01, ***p < 0.001. dDG, dorsal dentate gyrus; EYFP, enhanced yellow fluorescent protein; 4-OHT, 4-hydroxytamoxifen; DOX, doxycycline; Sac, sacrifice; WT, wild type.

**Figure S6.**
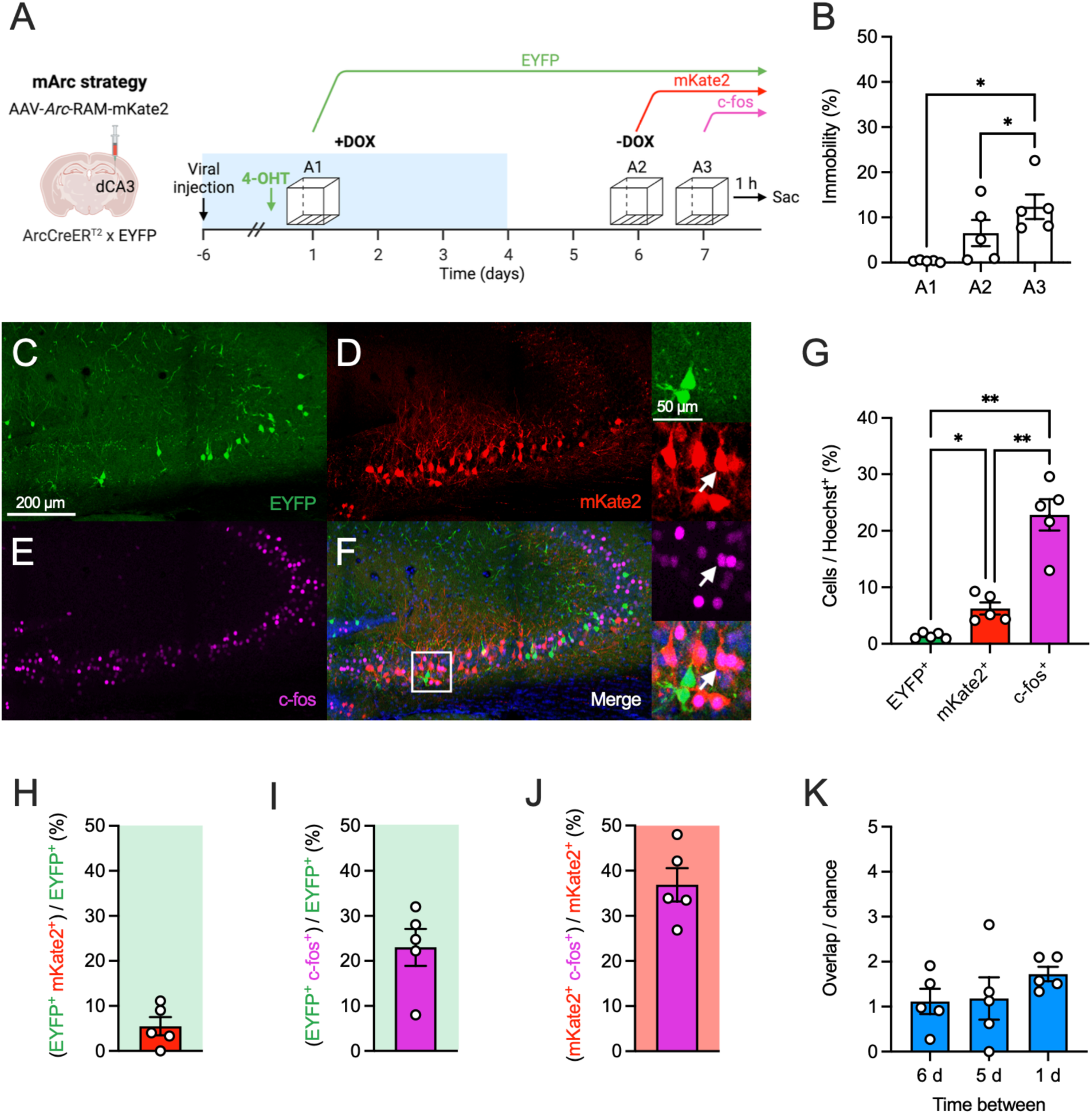
CA3 ensembles active during three exposures to the same context, related to Figure 4. (**A**) Experimental design to label CA3 ensembles active during three exposures to the same context. (**B**) Mice show increased immobility during the second and third exposures to context A compared to the first exposure. Representative image of: (**C**) EYFP^+^ cells, (**D**) mKate2^+^ cells, (**E**) c-fos^+^ cells, (**F**) merged CA3 section. The white box shows the area magnified in the right panels. The white arrow points to a mKate2^+^ c-fos^+^ cell. (**G**) The percentage of EYFP^+^, mKate2^+^ and c-fos^+^ CA3 cells. (**H**) The percentage of EYFP^+^ cells that also express mKate2 in CA3. (**I**) The percentage of EYFP^+^ cells that also express c-fos in CA3. (**J**) The percentage of mKate2^+^ cells that also express c-fos in CA3. **(K)** There is no difference in the overlap between ensembles active 6, 5, or 1 d apart in CA3. n = 5 mice. Error bars show mean ± SEM. *p < 0.05, **p < 0.01. dCA3, dorsal CA3; EYFP, enhanced yellow fluorescent protein; 4-OHT, 4-hydroxytamoxifen; DOX, doxycycline; Sac, sacrifice.

## SUPPLEMENTAL TABLES

**Table S1.** List of features for distinct contexts.

| **Feature** | **Context A** | **Context B** | **Context C** |
| --- | --- | --- | --- |
| Overall context | CFC chamber | CFC chamber | CFC chamber |
| Cleaning agent | Ethanol | Virkon-S | Sani-Cloth disposable wipes |
| Floor | Exposed steel bars | Exposed steel bars | Covered with plastic and bedding |
| Red lamps | Off | On | Off |
| Room lights | On | Off | On |
| Walls | None | Circular colored wall | Semi-circular colored wall |
| Scent | Lemon | Anise | None |
| Transfer cage | Standard home cage | White bucket | Standard home cage |
| Room | Room 1 | Room 1 | Room 2 |

**Table S2.** Antibody combinations for immunohistochemistry experiments.

| Figure Reference | Primary antibody, Company | Concentration | Secondary antibody, Company | Concentration |
| --- | --- | --- | --- | --- |
| Figure 1, S1 | Chicken anti-GFP, Abcam | 1:2,000 | Donkey anti-chicken Cy2, Jackson ImmunoResearch Labs | 1:500 |
|  | Rabbit anti-RFP, Rockland | 1:1,000 | Donkey anti-rabbit Cy3, Jackson ImmunoResearch Labs | 1:500 |
|  | Rat anti-c-fos, Synaptic Systems | 1:5,000 | Donkey anti-rat Alexa Fluor 647, Abcam | 1:500 |
| Figure 1, 2, 4, 5, 6, S3, S5, S6 | Chicken anti-GFP, Abcam | 1:2,000 | Donkey anti-chicken Cy2, Jackson ImmunoResearch Labs | 1:500 |
|  | Rat anti-c-fos, Synaptic Systems | 1:5,000 | Donkey anti-rat Alexa Fluor 647, Abcam | 1:500 |
| Figure S4 | Chicken anti-GFP, Abcam | 1:2,000 | Donkey anti-chicken Cy2, Jackson ImmunoResearch Labs | 1:500 |
|  | Rabbit anti-PV, Swant | 1:3,000 | Donkey anti-rabbit Alexa Fluor 647, Thermo Fisher Scientific | 1:500 |
| Figure S4 | Chicken anti-GFP, Abcam | 1:2,000 | Donkey anti-chicken Cy2, Jackson ImmunoResearch Labs | 1:500 |
|  | Rabbit anti-NeuN, Abcam | 1:500 | Donkey anti-rabbit Alexa Fluor 647, Thermo Fisher Scientific | 1:500 |

**Table S3.**
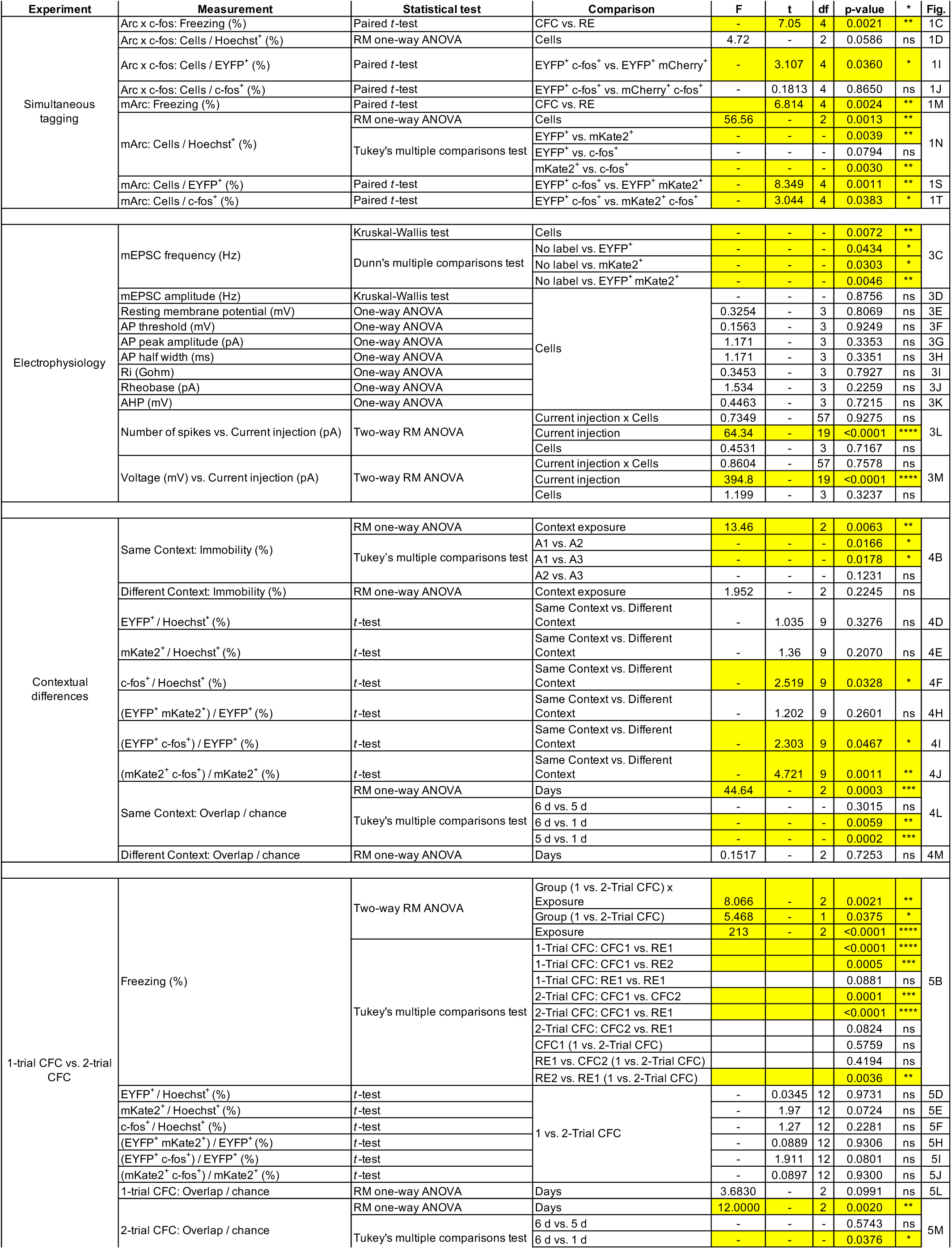
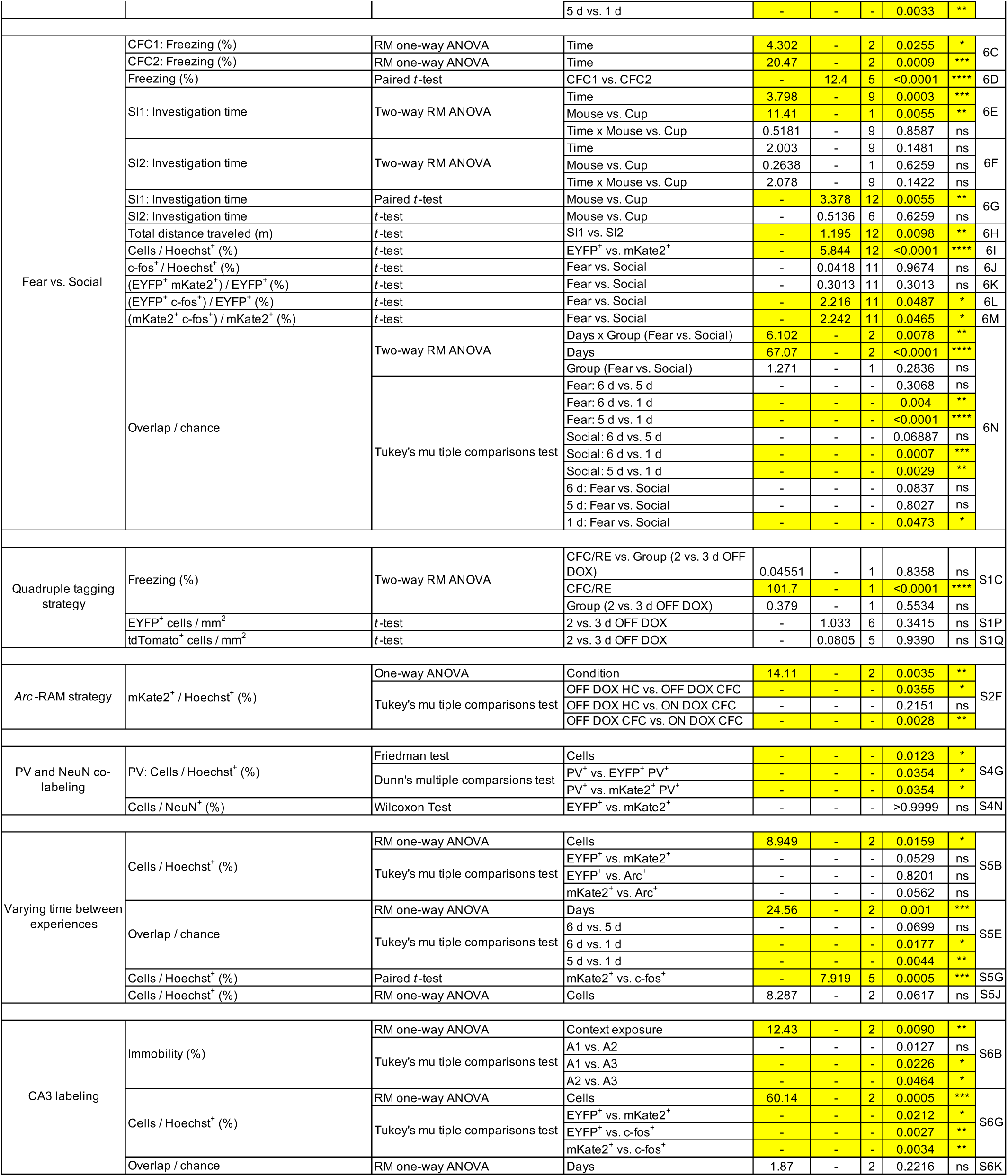
Statistical information.

